# Overcoming breeding constraints in polyploid oat from evolutionary insights

**DOI:** 10.1101/2025.07.18.665467

**Authors:** Xue Bai, Miao Sun, Yue Zhang, Hang Yin, Xue Cui, Xiaolin Liu, Zhigui Bao, Zhiyang Zhang, Zheng Dong, Zhongjie Liu, Yaoyao Wu, Dongwen Wang, Zhiwen Zou, Jia He, Mengting Li, Lihua Zheng, Yang Liu, Yao Zhou

**Author notes:** These authors contributed equally: Xue Bai, Miao Sun, Yue Zhang, Hang Yin, Xue Cui.

## Abstract

Polyploidy provides adaptive advantages in plants by buffering deleterious mutations^1,2^. While polyploidization can enhance agronomic traits such as increased biomass, known as the gigas effect^1–4^, increasing genetic gain in polyploid crops remains a critical but difficult goal due to the difficulty in dissecting complex trait inheritance^5,6^. Here we present chromosome-scale genome assemblies for 26 *Avena* taxa, spanning diploid, tetraploid, and hexaploid lineages. We traced four independent polyploidization events across the genus, including the formation of *A. agadiriana* as an allotetraploid (A_g_A_g_A_gʹ_A_gʹ_), and revealed a reticulate evolutionary history shaped by gene flow involving four subgenomes (A, B, C, D), for example, the hybrid speciation of *A. hirtula*. Transcriptomic analysis of 286 samples across 11 tissues, combined with deleterious mutation analysis from a cultivated population of 112 accessions, showed that polyploidization led to widespread functional redundancy among homoeologs, supporting a genome-wide buffering effect. However, derived allele frequency analysis revealed that, while disrupting functional genes may yield desirable traits, the buffering effect impedes the fixation of beneficial loss-of-function mutations and thereby limits breeding efficiency. Based on these integrative analyses, we propose breeding strategies to circumvent these limitations by targeting beneficial loss-of-function alleles within the complex polyploid background of oat. Our study highlights broader challenges in the improvement of polyploid crops and provides a foundation for future breeding strategies.

## Main

Polyploidy, the condition of possessing more than two complete sets of chromosomes, has long been recognized as a major force in plant evolution, driving species diversification and providing valuable genetic resources for breeding programs^1,3–5,7^. By introducing genome-wide redundancy, polyploidy not only enhances genetic diversity but also offers new opportunities for functional innovation, ultimately contributing to increased biological complexity^8,9^. Over the past few decades, extensive research has highlighted the benefits of polyploidy, such as gigas effect, buffering of deleterious mutations, and hybrid vigor, all of which have been effectively utilized in the development of many polyploid crops^2^. Nevertheless, the same gene redundancy that facilitates such innovation has also been described as a “geneticist’s nightmare”, as the presence of multiple gene copies with overlapping or compensatory functions can mask the phenotypic effects of natural mutations, reduce the predictability of genome editing outcomes, and hinder efforts to establish clear associations between genotype and phenotype^10^. Therefore, clarifying these genetic complexities is essential to fully harness the potential of polyploidy for breeding improvement.

Oat (*A. sativa*) is an important cereal crop and a major forage resource for livestock^11,12^. The genus *Avena* comprises up to 33 recognized species and/or subspecies across wild, weedy, and cultivated types, and it exhibits a well-established polyploid series that includes diploid (2n = 2x = 14), tetraploid (2n = 4x = 28), and hexaploid (2n = 6x = 42) cytotypes^13–16^. Among these, diploid species carry either the A or C genome, with the A genome further divided into subtypes such as A_a_, A_c_, A_d_, A_l_, and A_p_, while the C genome includes lineages such as C_v_ and C_e_. In contrast, tetraploid species predominantly exhibit AABB or CCDD genome compositions, with C-subgenome lineages including C_i_, C_g_, and C_y_, and D-subgenome lineages including D_i_, D_g_, and D_y_. Remarkably, all known hexaploid species share the AACCDD genome configuration, which originated through two successive allopolyploidization events^14,16,17^. This intricate variation in ploidy levels and subgenomic architectures, shaped by recurrent polyploidization and lineage divergence, positions *Avena* as an ideal model for investigating the evolutionary, genomic, and functional consequences of polyploidy in plants.

Here we present chromosome-scale *de novo* assemblies for 26 *Avena* species and/or subspecies, integrated with comprehensive transcriptomic profiling, to unravel how polyploidization reshapes gene expression dynamics and functional redundancy during *Avena* evolution. We demonstrate that polyploidization in oat confers a buffering effect against deleterious mutations; however, this mutational robustness comes at the cost of slowing the fixation of beneficial loss-of-function alleles. These findings reveal a trade-off between evolutionary stability and breeding efficiency in polyploid system, underscoring the complex consequences of polyploidization in shaping plant adaptation and improvement.

### Chromosome-level genome assembly reveals a new genome type in the *Avena* genus

To reconstruct the evolutionary history of *Avena* and evaluate the impact of polyploidization on oats, we collected a total of 26 species and/or subspecies within the genus, encompassing 12 diploids, 8 tetraploids, and 6 hexaploids, which represented nearly all known subgenome types (Extended Fig. 1a, Table S1 and Supplementary note 1). Using Pacific Biosciences (PacBio) high-fidelity (HiFi) reads at an average depth of 24.94× (from 21.75× to 28.71×), we generated *de novo* assemblies for all 26 taxa with contig *N_50_* sizes ranging from 69.77 Mb to 230.36 Mb (Fig. 1a and Table S2, 3), reflecting the high continuity of the assemblies. Chromosomal scaffolding and orientation were achieved using high-throughput chromatin conformation capture (Hi-C) reads (Table S4), which enabled the construction of 7, 14, and 21 pseudo-chromosomes for diploid, tetraploid, and hexaploid species, respectively. These pseudo-chromosomes were assigned to subgenomes according to the established oat nomenclature system^16^ (Supplementary note 2). We further evaluated the completeness of all assemblies using the k-mer (ranging from 97.76% to 98.93%) and BUSCO analysis (ranging from 99.06% to 99.49%) (Fig. 1b). The accuracy of all assemblies was assessed using QV value, all of which exceeded 69.86 (Fig. 1b). Together, these results highlight the exceptional quality of the genome assemblies.

**Fig. 1.**
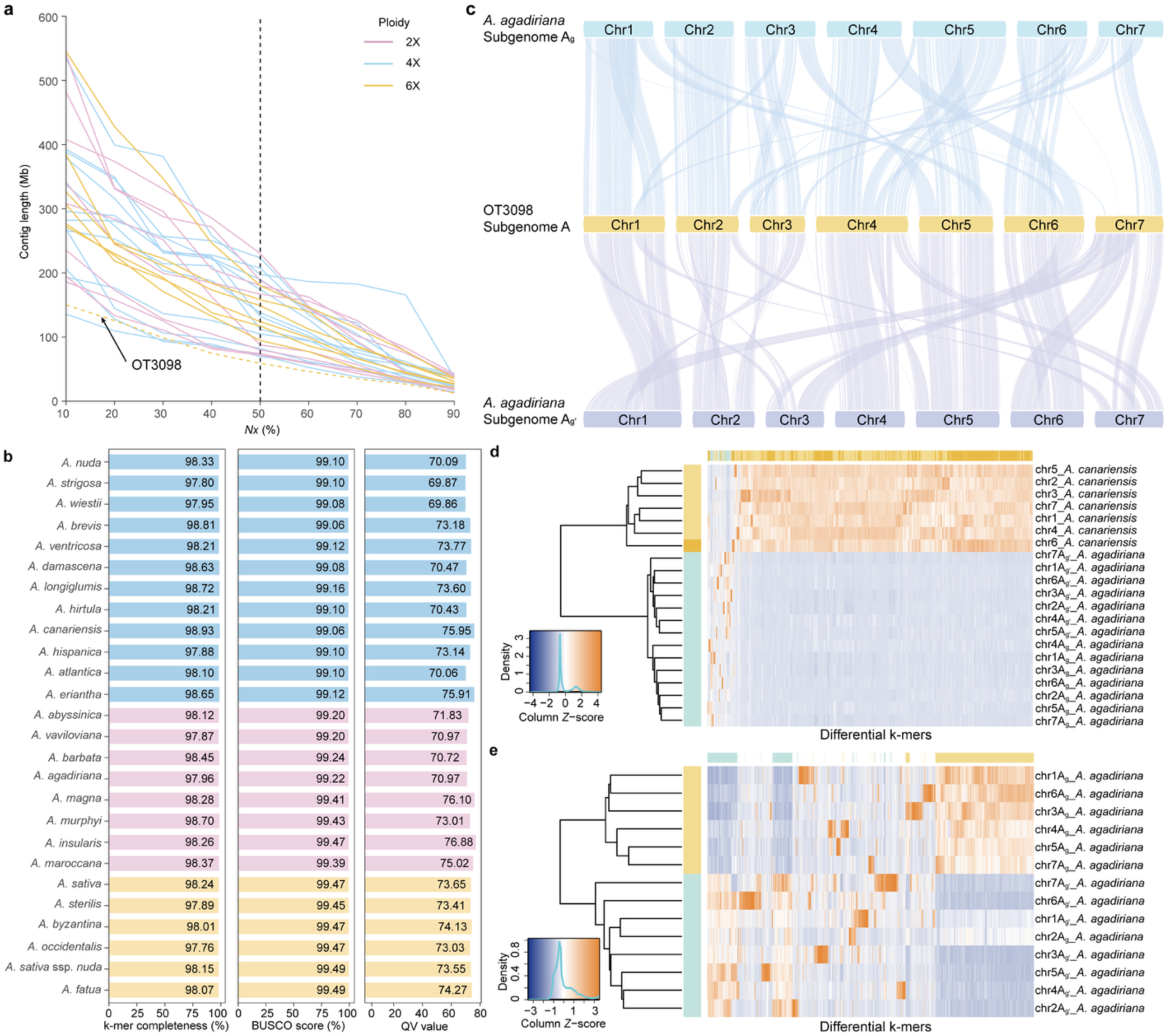
Genome assembly of the 26 *Avena* species and/or subspecies. **a**, Contig *Nx* sizes of all genome assemblies. Solid colored lines represent 26 *Avena* species and/or subspecies assemblies; the dotted line represents OT3098, indicated by the arrow. **b,** Completeness and accuracy evaluation for the 26 genome assemblies using k-mer completeness, Benchmarking Universal Single-Copy Orthologs (BUSCO), and QV value. **c,** Synteny among the subgenomes of *A. agadiriana* and the A subgenome of OT3098. **d, e,** Unsupervised hierarchical clustering revealed distinct clustering of the Ag and Agʹ subgenomes when *A. canariensis* (Ac) was included (**d**), whereas chr2 of *A. agadiriana* from the Ag subgenome clustered with the Agʹ chromosomes when *A. canariensis* was excluded (**e**).

Notably, the genome type of *A. agadiriana* remains controversial^17–20^. During the Hi-C scaffolding of *A. agadiriana*, we observed multiple contigs exhibiting high collinearity, which could not be reliably assigned to chromosomes, even after manual correction. To address this, we removed shorter duplicated contigs and re-scaffolded the remaining contigs using Hi-C data (Extended Fig. 1b and Supplementary note 3). Comparative genome distance analyses revealed that the two subgenomes of *A. agadiriana* are closely related to distinct subtypes of the A subgenome, designated as A_s_ and A_l_, while exhibiting greater divergence from the B, C, and D subgenomes (Extended Fig. 2). Based on these findings, we propose that the genome type of *A. agadiriana* be designated as A_g_A_g_A_gʹ_A_gʹ_. Further synteny analysis revealed extensive chromosomal fissions and fusions in the A_g_ and A_gʹ_ subgenomes compared to the A subgenome of OT3098 (https://wheat.pw.usda.gov/jb?data=/ggds/oat-ot3098v2-pepsico). For instance, chr3 of A_gʹ_ appears to have formed through the fusion of segments from chr1, chr2, chr6, and chr7, while the original chr3 was likely redistributed into chr1 and chr2 of A_gʹ_ (Fig. 1c). Intriguingly, SubPhaser analysis of k-mer patterns revealed that chromosomes from the A_g_ and A_gʹ_ subgenomes clustered separately when *A. canariensis* (A_c_ subtypes) was included (Fig. 1d). However, when *A. canariensis* was excluded, chr2 of A_g_ grouped with the A_gʹ_ chromosomes (Fig. 1e), suggesting that potential inter-subgenomic exchanges may have occurred between the A_g_ and A_gʹ_ subgenomes in *A. agadiriana*.

Following the assembly of all species and/or subspecies, we observed that the sizes of the C subgenomes (4.28 Gb on average) are visibly larger than those of the other three subgenomes (3.55 Gb for A, 3.48 Gb for B, and 3.31 Gb for D) (Table S5). Considering the gene numbers and sizes differ only slightly among the four subgenomes (Extended Fig. 3a and Table S6), we annotated the transposable elements (TE) across all subgenome assemblies to investigate the underlying genomic differences. TE content ranged from 83.91% to 86.67%, with long terminal repeat retrotransposons (LTR-RTs, 73.44%–76.98%) being the most abundant TE class. Among these, *Gypsy* LTR-RTs (46.75%–55.01%) were the most prevalent, followed by *Copia* LTR-RTs (18.40%–26.44%) (Table S7 and Supplementary note 4). The subgenome size exhibited a highly significant correlation (*r* = 0.998, *P* = 3.99 × 10^-53^, Wilcoxon test) with TE content (Extended Fig. 3b). A higher abundance of *Copia* LTR-RTs was found in C subgenomes (Extended Fig. 3c). In addition, insertion time estimates revealed two bursts of TE activity in the C subgenome, which likely contributed to its larger assembled size compared to the other subgenomes (Extended Fig. 3d). Therefore, the C subgenome likely underwent a distinct evolutionary trajectory characterized by extensive TE expansion.

### The phylogeny of the *Avena* subgenomes

To investigate the evolutionary history of *Avena* subgenomes and their respective subtypes (A, B, C, and D, with 8, 1, 6, and 4 subtypes in this study, respectively), we used a multispecies coalescent approach to infer a phylogeny based on 586 single-copy genes (SCGs), incorporating *O. sativa*, *B. distachyon*, *H. vulgare*, and *P. annua* as outgroup taxa. The resulting phylogeny delineated five monophyletic subgenome clades (Fig. 2a). The earliest diverged C clade encompassed all the six C subgenome subtypes (C_v_, C_e_, C_y_, C_g_, C_i_, and C_s_), followed by the emergence of the B clade, which retained only a single extant subtype. The A subgenome, the most structurally complex, consists of eight distinct subtypes and exhibits a paraphyletic structure, splitting into two sister clades (A1: A_g_, A_d_, A_c_; A2: A_a_, A_b_, A_gʹ_, A_l_, A_s_), with the intervening D clade (D_y_, D_g_, D_i_, and D_s_) positioned between them. Notably, the D clade shares the most recent common ancestor with the A2 lineage (Fig. 2a). These findings provide a phylogenetic basis for the historical misclassification of CCDD tetraploids as having an AACC genome composition^21^.

**Fig. 2.**
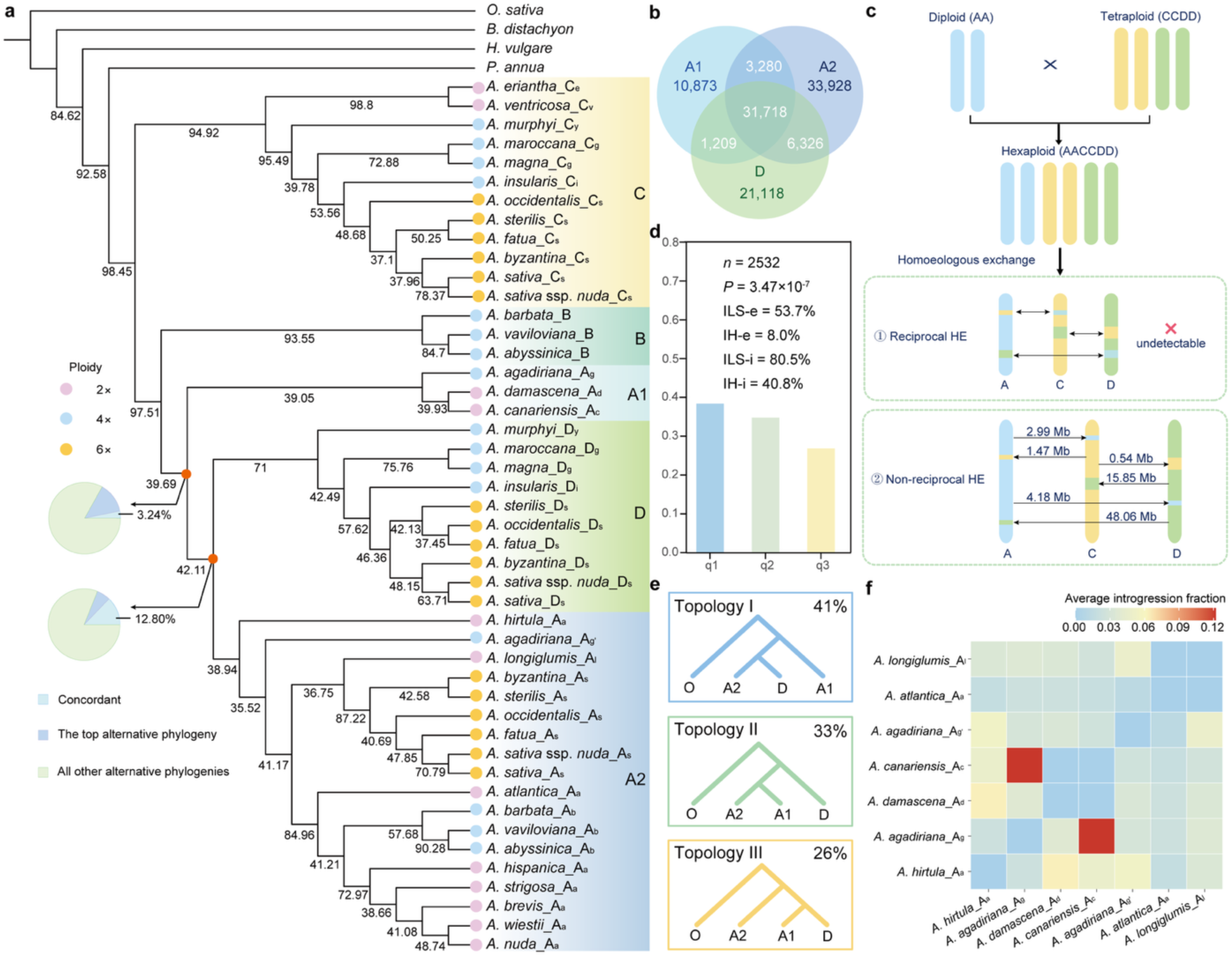
Subgenome origin of *Avena*. **a**, Phylogenetic relationships among the 46 oat subgenomes and four Poaceae outgroups. The coalescent-based tree, reconstructed from 586 single-copy genes (SCGs), groups the 46 oat subgenomes into five major clades (C, B, A1, D, and A2), distinguished by different background colors. The numbers on the nodes represent local posterior probability (localPP) values based on 586 individual gene trees. Pie charts on the nodes illustrate the proportion of gene trees supporting the common topology (blue), the primary alternative topology (green) and other alternative topologies (orange). Branch lengths are not scaled to reflect evolutionary distances. **b,** Venn diagram showing the numbers of shared and unique gene orthogroups in the A1, A2, and D clades. **c,** Illustration of homoeologous exchange (HE) in hexaploid oats. The numbers indicate the lengths of genomic segments derived from diploid and tetraploid ancestral homologous regions that exhibit signatures of non-reciprocal HE in hexaploids. Arrows indicate the direction of HEs. Due to limitations in both methodology and available datasets, reciprocal HEs remain undetectable. **d,** Detection of incomplete lineage sorting (ILS) and introgression/hybridization (IH) indices based on the phylogenetic topologies of the A1, D, and A2 clades across 586 SCGs phylogenetic tree. *N* denotes the number of simulated gene trees, while *P* represents the *p*-value from the *χ^2^*test assessing whether the counts of topologies q2 and q3 are equal, ILS-e and IH-e indicate the proportions of gene tree topological incongruence attributable to incomplete lineage sorting (ILS) and introgression/hybridization (IH), respectively. ILS-I and IH-I correspond to the calculated indices for ILS and IH. **e,** Proportions of gene tree topologies among the 586 genes concerning the topologies of A1, A2, and D clades. **f,** Heat map illustrating the average introgression fraction between each pair of subgenomes within the A1 and A2 clades. The A1 clade includes *A. agadiriana*_Ag, *A. damascene*_Ad, and *A. canariensis*_Ac, while the A2 clade comprises *A. hirtula*_Aa, *A. agadiriana*_Agʹ, *A. longiglumis*_Al, and *A. atlantica*_Aa.

However, extensive topological discordance was observed between individual gene trees and the coalescent-based phylogeny, particularly in the relationships among the A1, D, and A2 clades, which coincided with lower local posterior probability (localPP) values (localPP value = 42.11) (Fig. 2a). To reconcile these discrepancies, we employed the CASTER-pair^22^ method to reconstruct subgenomes relationships using the same set of 586 SCGs. In contrast to coalescent-based approach, CASTER-pair^22^ resolved the 46 *Avena* subgenomes into four monophyletic clades, with the A1 and A2 clades forming sister lineages that collectively share the most recent common ancestor with the D clade (Extended Fig. 4a). To systematically evaluate the relationship among A1, A2, and D clades, we conducted additional phylogenetic analyses using 2,889 SCGs identified across the A and D subgenomes, with the B subgenome designated as the outgroup. The incongruence persists (Extended Fig. 4b, c). Genome distance estimates further supported closer divergence between the A2 and D clades (0.04492) than between A1 and A2 clades (0.04619) (Extended Fig. 2). Additionally, the A2 clade shared more gene orthogroups with the D clade (6,326) than with the A1 clade (3,280) (Fig. 2b). The synonymous substitution rates (*Ks*) value between A2 and D clade (0.003) was lower than that between the A1 and A2 clade (0.018) (Extended Fig. 4d). Given the potential confounding effects of homoeologous exchange (HE) events in hexaploids on phylogenetic topology, we quantified genome-wide non-reciprocal HE frequencies (Supplementary note 5). Approximately 73.09 Mb of diploid and tetraploid ancestral homologous regions displayed non-reciprocal HE signatures in hexaploids (Fig. 2c). However, reciprocal HEs, which can also impact phylogenetic topology, remain undetectable with current dataset and methods. To minimize the influence of HE on phylogenetic topology, we excluded hexaploid individuals from the analysis, and recovered the same topological relationships among the A1, D, and A2 clades (Extended Fig. 4e). Collectively, these results suggest that while coalescent-based method likely capture deeper phylogenetic signals, persistent methodological discordance reflects evolutionary complexities within the *Avena* genus.

To further investigate the causes of discordance in the coalescent-based phylogenetic topology, we detected extensive signals of incomplete lineage sorting (ILS) and introgression (Extended Fig. 5a), particularly focusing on the topologies of A1, D, and A2 clades (Fig. 2d). We then examined the gene topologies within these clades, which exhibited an even genome-wide distribution (Extended Fig. 5b). The most frequent topology, representing 41% of ortholog groups, placed the D clade as sister to the A2 clade (Topology Ⅰ), consistent with the bifurcating tree structure. However, 33% of ortholog groups indicated that A2 was sister to A1 (Topology Ⅱ), a result that was significantly more frequent (*P* = 0.0105; *χ^2^*test) than the 26% observed for Topology Ⅲ, where A2 formed a distinct clade (Fig. 2e). The unequal distribution of these alternative topologies contradicts expectations under a model dominated solely by ILS^23^. QuIBL (Quantifying Introgression via Branch Length)^24^ analysis further indicated extensive gene flow within the A1 and A2 clades (Fig. 2f), suggesting that gene flow between these clades contributed to an increase in the proportion of Topology Ⅱ. Together, these results indicate that the evolutionary history of *Avena* subgenomes is shaped by a complex interplay of ILS and gene flow, leading to pervasive phylogenetic discordance.

### Reticulate evidence reveals hybrid origin and sequential polyploidization in oats

Although a well-resolved phylogenetic topology was obtained for oat subgenome lineages, the polytomy test revealed that the null hypothesis of hard polytomy could not be rejected at several shallow nodes in the phylogenetic tree, as the *P*-values are 0.51, 0.20, and 0.16, all greater than 0.05 (Extended Fig. 6a). This suggests unresolved phylogenetic relationships within these lineages. Specifically, discrepancies observed among the A-genome diploids, accompanied by lower localPP values, cannot be solely attributed to ILS (Fig. 2a and Extended Fig. 6b). This observation supports a reticulate evolutionary scenario among the A-genome diploids species and/or subspecies. Both QuIBL^24^ and network analyses revealed substantial ancient gene flow events (Fig. 2f and Extended Fig. 7a), particularly involving *A. hirtula*. Notably, the phylogenetic position of *A. hirtula* shifted from the A2 to the A1 clade when 2,889 single-copy orthologs were used to reconstruct the phylogenetic tree (Extended Fig. 4b). To investigate whether ILS is the main cause of the observed phylogenetic patterns, we simulated the expected distribution of gene tree topologies under an ILS-only scenario. For each gene sequence type analyzed, we observed that the ratios of Tree-II and Tree-III topologies differed significantly (*P* = 1.658 × 10^-8^, two-tailed Student’s *t*-test) from those predicted by the ILS-only simulations (Extended Fig. 7b). This discrepancy led us to reject the hypothesis that ILS alone accounts for the phylogenetic discordance. Instead, it supports the hypothesis of a hybrid origin for *A. hirtula*. Furthermore, PhyloNet^25^ analysis suggests that the *A. hirtula* may originate from an ancient hybridization event between the ancestor of *A. damascene* and the common ancestor of *A. atlantica* and *A. longiglumis* (Extended Fig. 7c–g). Similar evolutionary complexity was observed in hexaploids, where conflicting topologies among A, C, and D subgenomes (Fig. 2a), corroborated by QuIBL^24^ analysis (Fig. 3a–c), indicate extensive gene flow within these subgenomes. Crucially, the patterns of gene flow exhibited significant subgenome-specific variation (Fig. 3a-c), implying distinct evolutionary trajectories for each subgenome within the hexaploid oats. These findings highlight that, despite genome-scale data, extensive reticulate evolution and lineage-specific gene flow continue to obscure the phylogenetic resolution within several *Avena* species and/or subspecies, preventing the reconstruction of a fully resolved evolutionary history in this study.

**Fig. 3.**
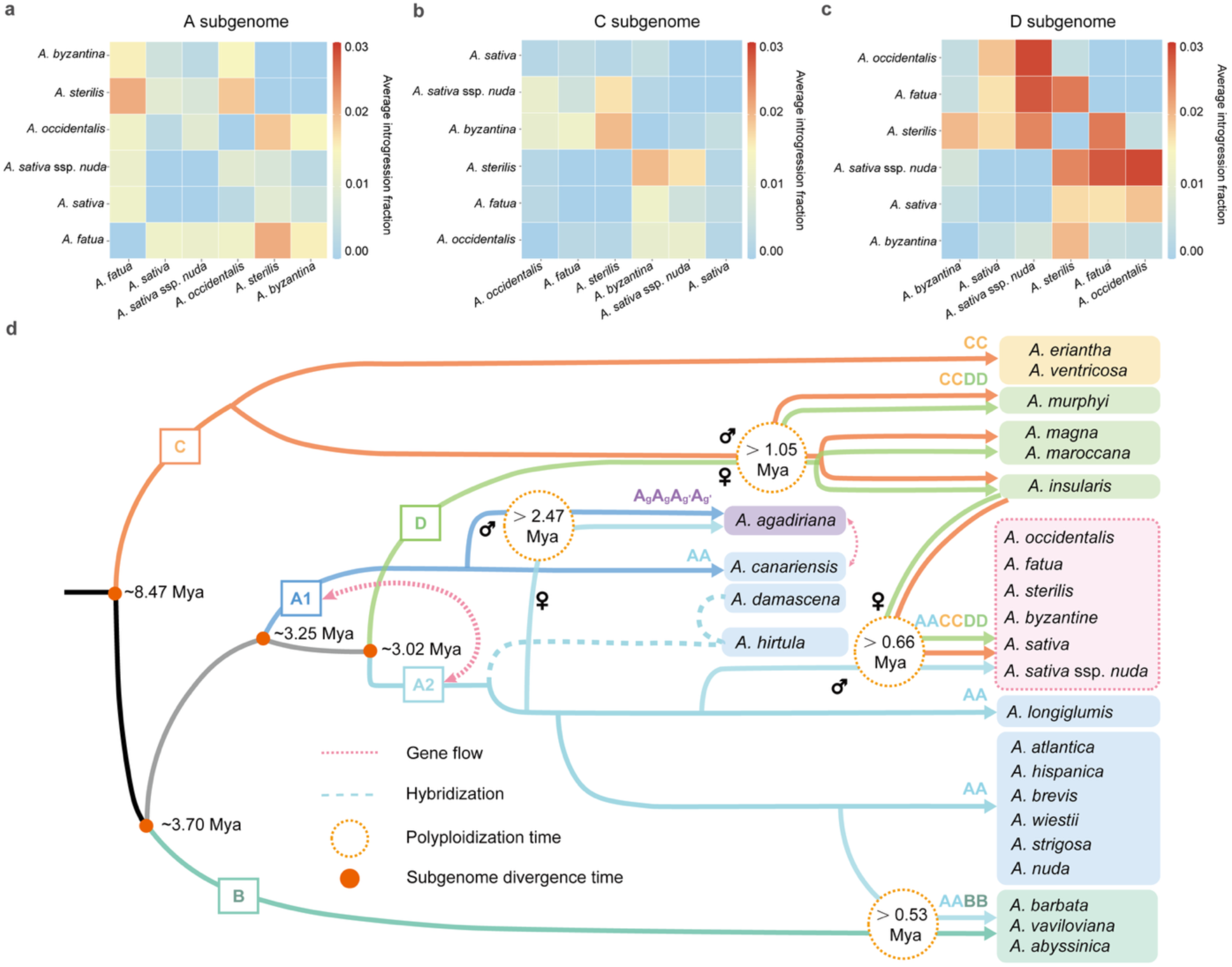
Reticulate evolution and polyploidization of *Avena*. **a–c**, Heat maps showing the average introgression fractions between each pair of A subgenomes (**a**), C subgenomes (**b**), and D subgenomes (**c**) within hexaploid oats. **d,** Model depicting the origins and evolutionary history of oat subgenomes and the polyploidization events leading to the formation of tetraploid and hexaploid species and/or subspecies. The five subgenome types (A1, A2, B, C, and D) are represented in distinct colors. Approximate dates for subgenome divergence (million years ago, Mya) are indicated beside red points. Approximate dates for polyploidization events are indicated within circles. Potential gene flow and hybridization event are represented by red and blue dashed lines, respectively.

We further assembled chloroplast genomes from 26 *Avena* species and/or subspecies to investigate the maternal genome donor of polyploid oats (Extended Fig. 7h and Supplementary note 6). The results indicated that hybridization between the diploid progenitors of A_g_ and A_gʹ_ followed by polyploidization, gave rise to *A. agadiriana* (A_g_A_g_A_gʹ_A_gʹ_), with the progenitor of A_gʹ_ potentially serving as the maternal donor. For AABB-genome tetraploids, A_a_ of *A. atlantica* is the closest relative to the A subgenome of tetraploid oats in the SCG-based phylogenetic tree. Additionally, the C_v_ or C_e_ diploids, which diverged earliest, occupy distinct subclades separate from the CCDD-genome tetraploids. This suggests they may not be the C-genome donors, leaving the ancestral C-genome contributor to CCDD tetraploids unresolved. The hexaploid oat (AACCDD) originated through hybridization between a paternal Al-genome diploid ancestor, closely related to the progenitor of *A. longiglumis*, and a maternal CCDD-genome tetraploid oat, closely related to *A. insularis*. However, the B-genome, as well as the D-genome diploid progenitors remain unidentified and are likely extinct. Moreover, divergence time was estimated via secondary calibrated molecular clocks (Supplementary note 7), revealing that the C and B/A1/D/A2 lineages diverged before ∼8.47 Mya (5.27–11.09 Mya, with 95% confidence interval, CI), followed by the divergence of the B and A1/D/A2 lineages before ∼3.70 Mya (2.34–4.72 Mya, with 95% CI). Subsequently, the A1 and D/A2 lineages separated before ∼3.25 Mya (2.05–4.13 Mya, with 95% CI), and finally, the D and A2 lineages diverged before ∼3.02 Mya (1.91–3.85 Mya, with 95% CI). In addition, all four polyploidization events in *Avena* genus occurred during the Pleistocene. The earliest, dating to no later than ∼2.47 Mya, gave rise to the A_g_A_g_A_gʹ_A_gʹ_ tetraploid. The second event, before ∼1.05 Mya, led to the formation of the CCDD-genome tetraploid. The third and fourth polyploidization events, estimated no later than ∼0.53 Mya and ∼0.66 Mya, respectively, led to the formation of the AABB-genome tetraploid and the AACCDD hexaploid.

Together, we proposed a refined model for the origins and polyploidization events within the *Avena* genus (Fig. 3d), highlighting the complex polyploidization and reticulate evolution that shaped the evolutionary history of this genus.

### Balanced and divergence gene expression following oat polyploidization

Polyploidization can result in subgenome bias, where one subgenome becomes dominant^26^, and may also alter gene expression profiles^27^. To explore its impact in oats, we analyzed 286 RNA-seq samples from the 26 *Avena* taxa, covering 11 tissue types from leaf, root, panicle, and grain at four developmental stages (Fig. 4a). Our analysis revealed a U-shaped expression distribution across tissues (Fig. 4b), mirroring patterns observed in humans^28^. Moreover, a substantial proportion of genes were either constitutively expressed across all 11 tissues (ranging from 9.75% to 27.45%, with an average of 18.34%) or completely unexpressed (ranging from 31.10% to 47.18%, with an average of 39.32%), and this pattern was consistent across subgenomes in both tetraploids and hexaploids (Fig. 4b and Fig. S1). Notably, consistent with previous findings in human^28^, transcription within each tissue was driven by a relatively small subset of highly expressed genes, with the top 1,000 genes accounting for approximately 50% of total expression (Fig. 4c). When focusing on these top 1,000 genes—identified in at least one tissue across the 26 *Avena* taxa—we found no evidence of subgenome bias in their expression (*P* = 1, proportion *z*-test) (Table S8). However, the proportion of non-expressed genes increased in tetraploids (40.38%) and hexaploids (45.59%) relative to diploids (35.57%), whereas the proportion of constitutive genes declined (22.34%, 15.68%, and 13.63% in diploids, tetraploids, and hexaploids respectively) (Extended Fig. 8a), suggesting that polyploidization alters gene expression profiles in oats to some extent.

**Fig. 4.**
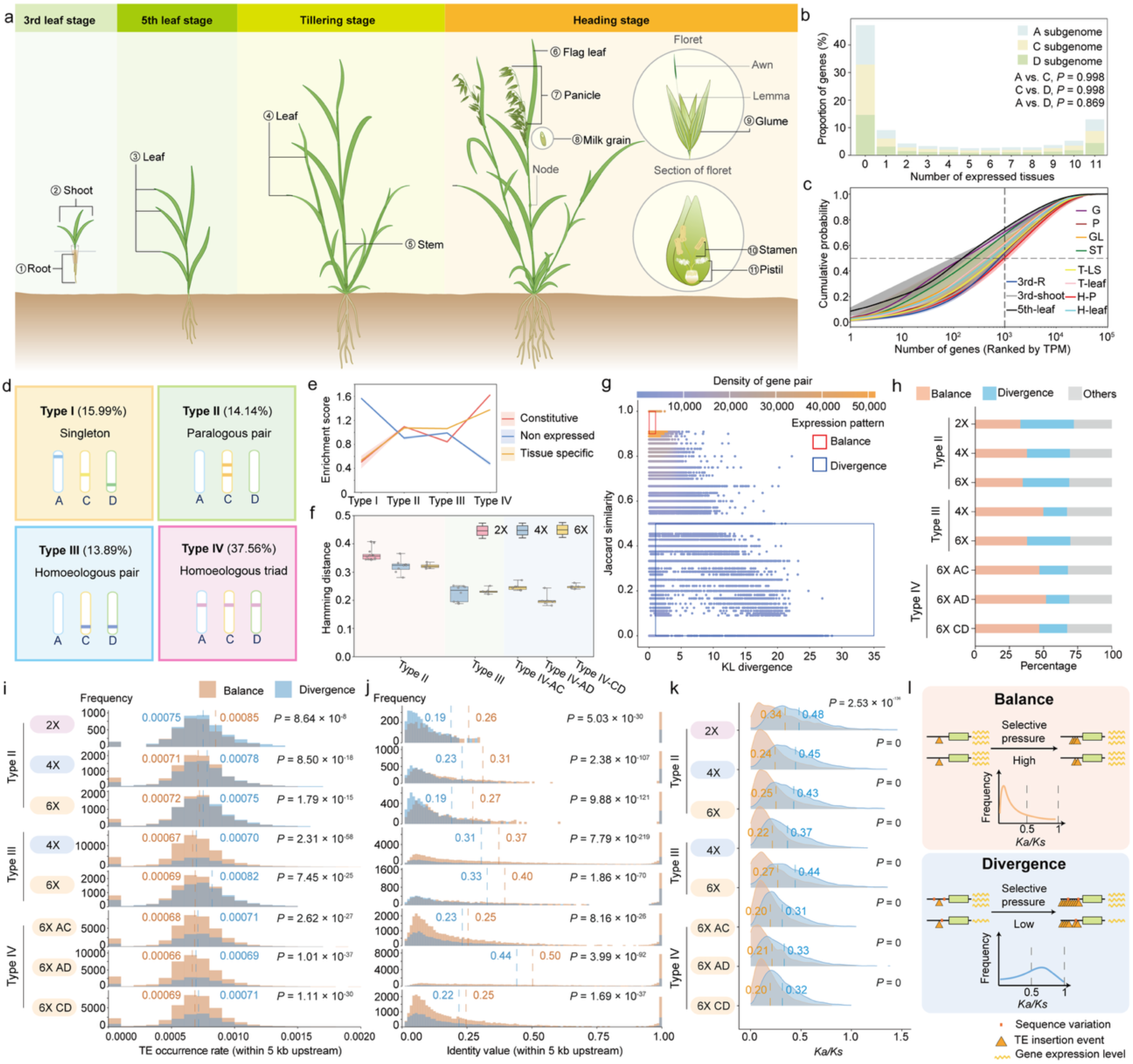
Polyploidization in *Avena* preserves widespread gene expression balance among homologous genes. **a**, Overview of tissues sampled for transcriptomic analysis across multiple growth stages. Tissues labeled 1–11 represent those used for RNA sequencing. **b,** Proportion of genes expressed across different numbers of tissues in *A. sativa* ssp. *nuda*. Genes with transcripts per million (TPM) > 1 were considered expressed. The proportion of genes expressed in the same number of tissues showed no significant differences among A, C, and D subgenomes (Kolmogorov-Smirnov test, *P* > 0.05). **c,** Cumulative distribution of the mean fractional contribution of gene expression levels to total transcription across tissues. Genes are ranked by TPM from highest to lowest within each tissue (x-axis). Curves represent mean values across 26 *Avena* species and/or subspecies for the corresponding tissue, with lighter-colored surfaces around the mean representing the standard error. 3rd-R and 3rd-shoot, root and shoot at 3rd leaf stage. 5th-leaf, leaf at 5th leaf stage. G, milk grain. P, pistil. GL, glume. ST, stamen. T-LS and T-leaf, leaf sheath and leaf at tilling stage. H-P and H-leaf, panicle and flag leaf at heading stage. **d,** Classification of genes based on copy number and their distribution among subgenome. Percentage values indicate the proportion of each gene category relative to the total gene count in hexaploids. **e,** Enrichment scores for gene types I–IV corresponding to distinct expression profiles in hexaploids. Constitutive: gene expressed across all 11 tissues. Tissue-specific: gene expressed in 1-10 tissues. Non-expressed: gene not expressed in any of the 11 tissues. The lighter-colored surfaces around the mean represent dispersion, calculated as the standard deviation divided by the mean. **f,** Expression level differences between gene pairs in Types II–IV, evaluated using Hamming distance. Ploidy levels are color-coded: 2× (pink), 4× (blue), and 6× (yellow). **g,** Distribution of Homoeologous traid (A–C subgenomes) of hexaploids based on their Kullback-Leibler (KL) divergence and Jaccard similarity in expression. Two expression groups were identified: Balance, defined by KL divergence in the range [0, 1] and Jaccard similarity in the range [0.9, 1.0]; and Divergence, defined by KL divergence in the range [1, 35] and Jaccard similarity in the range [0, 0.5]. **h,** Proportion of gene pairs in Types II–IV, classified expression patterns as Balance, Divergence, or Other. **i, j,** TE occurrence rate (**i**) and identity value (**j**) of balanced and divergent genes in Types II–IV, analyzed using 5-kb 5’ upstream regions. Dashed lines indicate mean values. **k,** Comparison of *Ka*/*Ks* between balanced and divergent genes in Types II–IV. Dashed lines indicate mean values. **i–k,** *P* values were calculated using Wilcoxon test. **l,** Schematic illustration of the potential mechanism underlying gene pair expression balance/divergence and TE accumulation. In the Balance scenario (top), high identity in upstream regions maintains similar gene expression levels between gene pairs. Strong selective pressure, reflected by lower *Ka*/*Ks* ratios, restricts transposable element (TE) insertions. In contrast, in the Divergence scenario (bottom), lower upstream identity leads to expression divergence. Relaxed selective pressure, indicated by higher *Ka*/*Ks* ratios, allows greater TE accumulation. Symbols represent gene pairs (green rectangles), sequence variation (orange squares), TE insertions (orange triangles), and gene expression levels (orange waves).

To further investigate gene expression dynamics following polyploidization, we examined genes with varying copy numbers (Fig. 4d). Among the 26 *Avena* taxa, single-copy genes (singletons) made up 65.89% of the total gene count in diploids, but this proportion decreased to 18.98% in tetraploids and 15.99% in hexaploids (Extended Fig. 8b). In contrast, the proportion of two-copy and multi-copy genes increased with polyploidy (Extended Fig. 8b). Interestingly, in hexaploids, singletons (Type I) were more likely to be non-expressed (enrichment score = 1.57), whereas homoeologous triads (Type IV) tended to be constitutively expressed (enrichment score = 1.64) (Fig. 4e). Specifically, 71.48% of singletons were non-expressed, whereas this proportion dropped to only 21.73% for homoeologous triads. To further assess divergence in expression profiles among homologous gene pairs, we calculated Hamming distances based on binary expression (expressed vs. non-expressed) across tissues. The results indicated that expression divergence was lower among homoeologs than among paralogs. In addition, within homoeologous triads, the A and D subgenomes were more similar to each other (AD = 0.197) than either was to the C subgenome (AC = 0.246, DC = 0.249), suggesting that the earlier a homolog originated, the more likely its expression profile has diverged over time (Fig. 4f). Furthermore, to evaluate divergence in tissue specificity and expression levels among homologs, we applied Kullback– Leibler (KL) divergence and Jaccard similarity analyses across gene categories (Types II–IV) and ploidy levels (Fig. 4g and Extended Fig. 8c–i). The results revealed that balanced homoeologs accounted for a higher proportion than those with divergent expression in polyploids (averaging 48.41% vs. 19.46%), with this trend being particularly pronounced in homoeologous triads of hexaploids (Fig. 4h). In all six hexaploids, 55,905 (54.17%) homoeologous triads exhibited balanced expression in at least one subgenome pair, with 37,374, 42,647, and 37,109 gene pairs showing balanced expression between the A–C, A–D, and C–D subgenomes, respectively. Together, these findings suggest that while gene expression divergence occurs across all gene types, homoeologous genes are more likely to maintain balanced expression following polyploidization in oats, compared to singletons or paralogs.

To explore the mechanisms underlying balanced versus divergent gene expression, we quantified differences in TE content and sequence identity within the 5- and 50-kb 5’ upstream regions of both balanced and divergent gene pairs. Specifically, TE occurrence rates and sequence identity in these upstream regions were assessed, and results showed that TE occurrence rates were significantly higher between divergent genes, while sequence identity was significantly higher between balanced ones (Fig. 4i, j and Extended Fig. 9a, b; Wilcoxon test). Nevertheless, divergent genes exhibited significantly higher *Ka*/*Ks* ratios than balanced genes (Fig. 4k; Wilcoxon test), indicating that they are under weaker purifying selection. This observation aligns with previous findings that TEs preferentially insert into genomic regions subject to reduced selective pressure^29^. Therefore, we propose that upstream sequence variation—rather than TE content alone—plays a more critical role in driving expression divergence. Furthermore, relaxed selective pressure likely facilitates the accumulation of TEs in the 5’ upstream regions of divergent genes, which may, in turn, reinforce gene expression divergence (Fig. 4l).

Overall, these findings highlight the complex interplay between genome duplication, regulatory architecture, and evolutionary pressures in shaping gene expression patterns across ploidy levels. In *Avena*, polyploidization does not lead to subgenome dominance but instead maintains widespread gene expression balance among homologous genes, particularly in hexaploid oats.

### Contrasting effect of polyploidization on hexaploid oat breeding

Despite the widespread balance in gene expression, we further investigated potential functional redundancy among homologs in hexaploid oat by reanalyzing whole-genome sequencing data from a diverse panel of 112 cultivated accessions^30^, using *A. sativa* ssp. *nuda* (AACCDD) as the reference genome. In total, we identified 67,007,966 SNPs and 3,856,289 indels. Genome-wide estimation of nucleotide diversity (*π*) revealed that the C and D subgenomes harbored comparable levels (𝜋_*c*_= 1.27 × 10^-^³, 𝜋_*D*_ = 1.34 × 10^-^³) of genetic diversity, both of which were higher than that observed in the A subgenome (𝜋_*A*_= 8.24 × 10^-^⁴). Moreover, homoeologous triads, particularly those with balanced expression, exhibited the lowest nucleotide diversity (*π* = 2.46 × 10^-4^, balanced homoeologous triads vs. balanced homoeologous pairs, *P* = 3.71 × 10^-29^, Wilcoxon test) (Fig. 5a), suggesting that these genes are subject to purifying selection, consistent with the *Ka*/*Ks* patterns previously observed (Fig. 4k). Notably, non-expressed homoeologous triads displayed similar nucleotide diversity (*π* = 4.21 × 10^-4^) to that of balanced paralogous pairs (*π* = 4.67 × 10^-4^) (Fig. 5a), hinting that these genes may have experienced purifying selection prior to polyploidization. In addition, nucleotide diversity comparisons across homologs showed that over 26.51% of balanced gene pairs exhibited more than a 10-fold difference in diversity, a disparity comparable to that observed in divergent gene pairs (Fig. 5b), implying that balanced homologs, despite similar expression levels, may have undergone functional divergence.

**Fig. 5.**
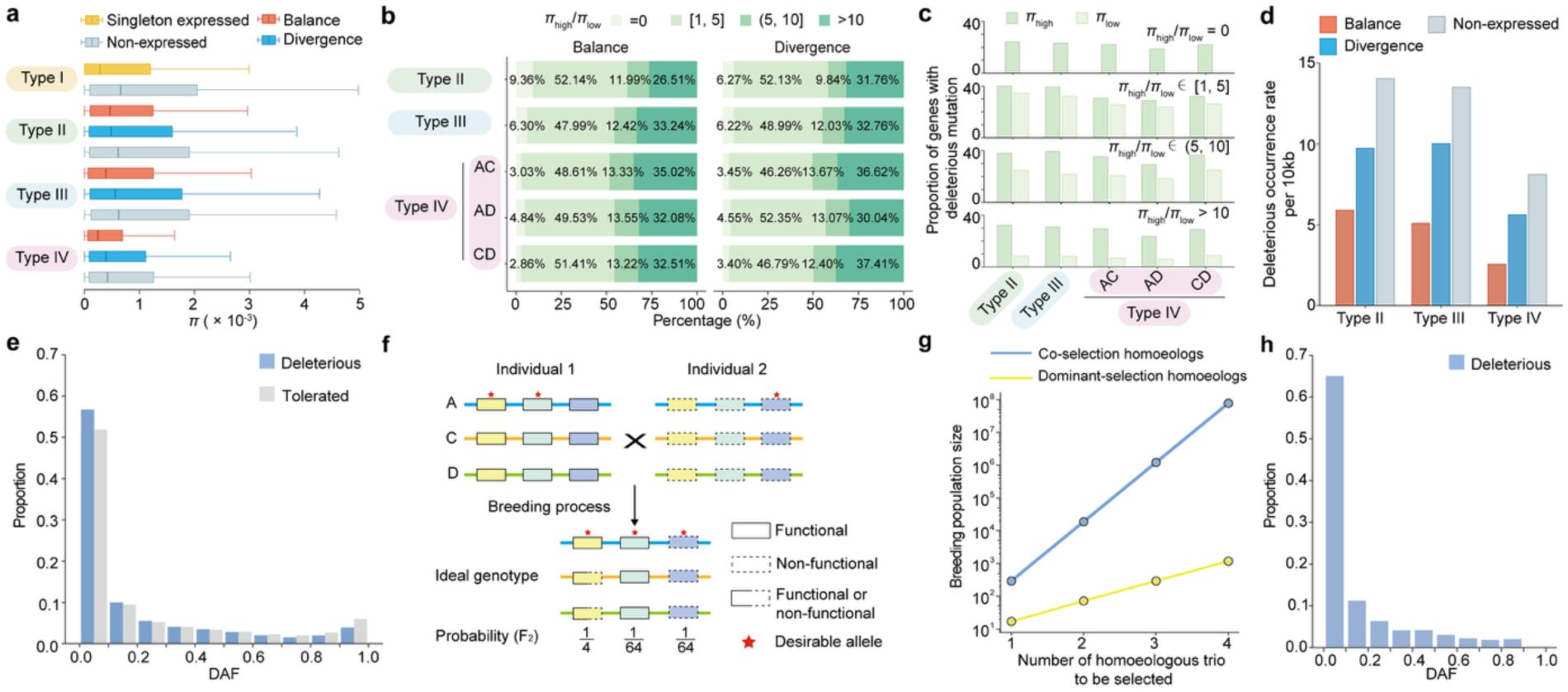
Polyploid buffering limits the fixation of beneficial deleterious mutations in breeding. **a**, Nucleotide diversity (*π*) across different gene types in four gene categories (Type I–IV). **b,** Distribution of nucleotide diversity ratio (*π*high/*π*low) among gene pairs in Balance and Divergence categories across gene types II–IV. Gene pairs were grouped into four ratio intervals: 0, [1–5], (5–10], and >10. Type IV genes are further classified into subgenomes AC, AD, and CD. **c,** Proportion of deleterious mutations in each homolog of balanced gene pairs. Balanced gene pairs were grouped based on their nucleotide diversity ratio (*π*high/*π*low). Analyses were conducted across gene types II–IV, with type IV further split into subgenomes AC, AD, and CD. **d,** Occurrence rate of deleterious mutations (per 10 kb) among genes classified as Balance, Divergence, and Non-expressed across gene types II–IV. **e,** Derived allele frequency (DAF) spectrum of deleterious (blue) and tolerated (gray) mutations. **f,** Schematic illustration of the probability of fixing desirable alleles in different genetic contexts. The diagram illustrates a breeding scenario where individuals carry different combinations of functional (solid lines) and non-functional (dashed lines) homoeologs. To obtain the ideal genotype in the F2 generation, where all three homoeologs carry either a desirable but deleterious allele (blue rectangles with dashed lines) or a dosage-dependent desirable allele (green rectangles with slide lines), the probability is only 1/64. In contrast, when selection targets a dominant homoeolog (red rectangles with slide lines), the probability of fixation increases significantly to 1/4. **g,** Population size requirements under different selection strategies. Scenarios compared include Co-selection (blue, selecting homoeologous traid simultaneously) and Dominant-selection (yellow, selecting only one functional copy), with population size increasing exponentially as the number of selected triads increases. **h,** Derived allele frequency (DAF) of deleterious mutations in the remaining gene copy of homologous gene pairs in which one copy harbors a high-frequency deleterious mutation (DAF > 0.9).

To explore the consequences of such divergence, we employed SIFT (Sorting Intolerant From Tolerant)^31^ scores and identified 98,612 deleterious mutations across 38,747 genes. Within balanced gene pairs, the copy with higher nucleotide diversity generally tend to harbor more deleterious mutations, and greater differences in nucleotide diversity between gene copies corresponded to more substantial differences in the number of deleterious variants (Fig. 5c). Additionally, balanced genes exhibited lower deleterious mutation occurrence rates than divergent and non-expressed genes, with triads showing the least accumulation (Fig. 5d), indicating that balanced genes are under positive selection—consistent with patterns observed in *Ka*/*Ks* ratios and 5’ upstream sequence conservation.

To assess the selection dynamics of deleterious mutation, we analyzed the derived allele frequency (DAF) distribution. A majority of deleterious mutations (55,964, 56.75%) were rare (DAF < 0.1), exceeding the corresponding proportion for tolerated mutations (257,265, 51.87%, *P* = 2.26 × 10^-10^, Wilcoxon test), suggesting an enrichment of rare deleterious alleles and ongoing purifying selection. In contrast, a fractional proportion (3,828, 3.88%) displayed high frequencies (DAF > 0.9) (Fig. 5e). Further analysis of nucleotide diversity for genes with deleterious mutations and those with DAF > 0.9 revealed that the latter had significantly lower nucleotide diversity (1.89 × 10^-^³ vs. 1.19 × 10^-^³, *P* = 0, Wilcoxon test), implying that some high-DAF loss-of-function mutations may be desirable and subject to positive selection during oat breeding. However, given the buffering effect among homoeologous gene copies, achieving fixation of a desirable loss-of-function mutation within a homoeologous triad would require deleterious mutations in all three copies (i.e., co-selection), drastically reducing the probability of obtaining the desired trait compared to targeting dominant-selection homoeologs (Fig. 5f). Consequently, this constraint imposes a significant challenge for breeding, as the population size needed to capture optimal combinations increases exponentially with the number of target loci (Fig. 5g). For instance, identifying one ideal individual with 99% probability in an F₂ population would require 293 individuals for a single triad but over 77 million for four triads—an unfeasible scale for breeding programs.

The situation becomes even more challenging when desirable loss-of-function genes do not carry any natural deleterious mutations. Among the 2,598 genes containing high-frequency deleterious mutations (DAF > 0.9), 266 (10.24%) were singletons, 255 (9.82%) were in balanced homologous gene pairs, and 398 (15.33%) were in divergent pairs. In cases where one gene copy within a homologous pair harbored a high-frequency deleterious allele, we observed that the corresponding homologs were largely unaffected, with 51.84% lacking any deleterious mutation. For homologs carrying such variants, the DAF between 0.4 and 0.5 was comparable to those between 0.3 and 0.4 (Fig. 5h), in contrast to the decreasing trend observed in all deleterious mutation (Fig. 5e). This pattern suggests that these loci may be undergoing positive selection but are subject to the decelerating effects of polyploid buffering. Taken the leaf size as an example, a key trait for both productivity and biomass^32^, we compiled a list of 124 genes previously reported to regulate leaf development in Arabidopsis, rice, wheat, and barley, and identified orthologs for 75 in *A. sativa* ssp. *nuda*. In total, 337 orthologs were obtained, with 99.71% retaining at least two homologs (Extended Fig. 10a). Expression pattern analyses revealed that 97.6% of these genes maintained balanced expression across leaf samples from different developmental stages (Extended Fig. 10b). Among the 75 orthologs, 58 were positive regulators and 17 were negative regulators. Importantly, no homoeologous group was found in which all copies harbored deleterious mutations (Table S9). In breeding, this redundancy supports selection for enhanced traits through positive regulators, but poses a challenge when targeting negative regulators, as all copies must be simultaneously disrupted to achieve a phenotypic effect.

Collectively, these findings highlight the contrasting role of polyploidy in oat breeding. On one hand, homoeolog redundancy serves as a genetic buffer that protects against deleterious mutations and contributes to trait stability across generations. On the other hand, this same redundancy complicates the selection and fixation of beneficial loss-of-function mutations, particularly when targeting negative regulators. As a result, breeding strategies in polyploid crops like oat must account for the complexity of gene copy number and the functional interplay among homoeologs to effectively harness genetic variation for crop improvement.

### Targeted breeding for hexaploid oat

Through an evolutionary dissection of how polyploidization reshapes gene expression dynamics and functional redundancy, we found that the intrinsic buffering and functional interplay among homologs impose substantial constraints on the efficiency of conventional breeding approaches. To overcome these limitations, we explored strategies to effectively harness beneficial loss-of-function genes and enhance trait improvement within the complex polyploid context of hexaploid oat (Fig. 6). A crucial first step is gene classification through comprehensive genomic screening to identify singletons, divergent homologs, partially redundant homologs, and fully redundant homologous triads. Genome-wide analyses based on redundancy levels, expression patterns, and functional divergence provide a foundation for designing breeding interventions tailored to the genetic architecture of each gene type. Building on this classification, high-frequency deleterious alleles that exhibit signs of positive selection emerge as promising targets for trait improvement. These alleles can be prioritized for direct selection or minimal genome editing, minimizing the reliance on extensive genetic engineering.

**Fig. 6.**
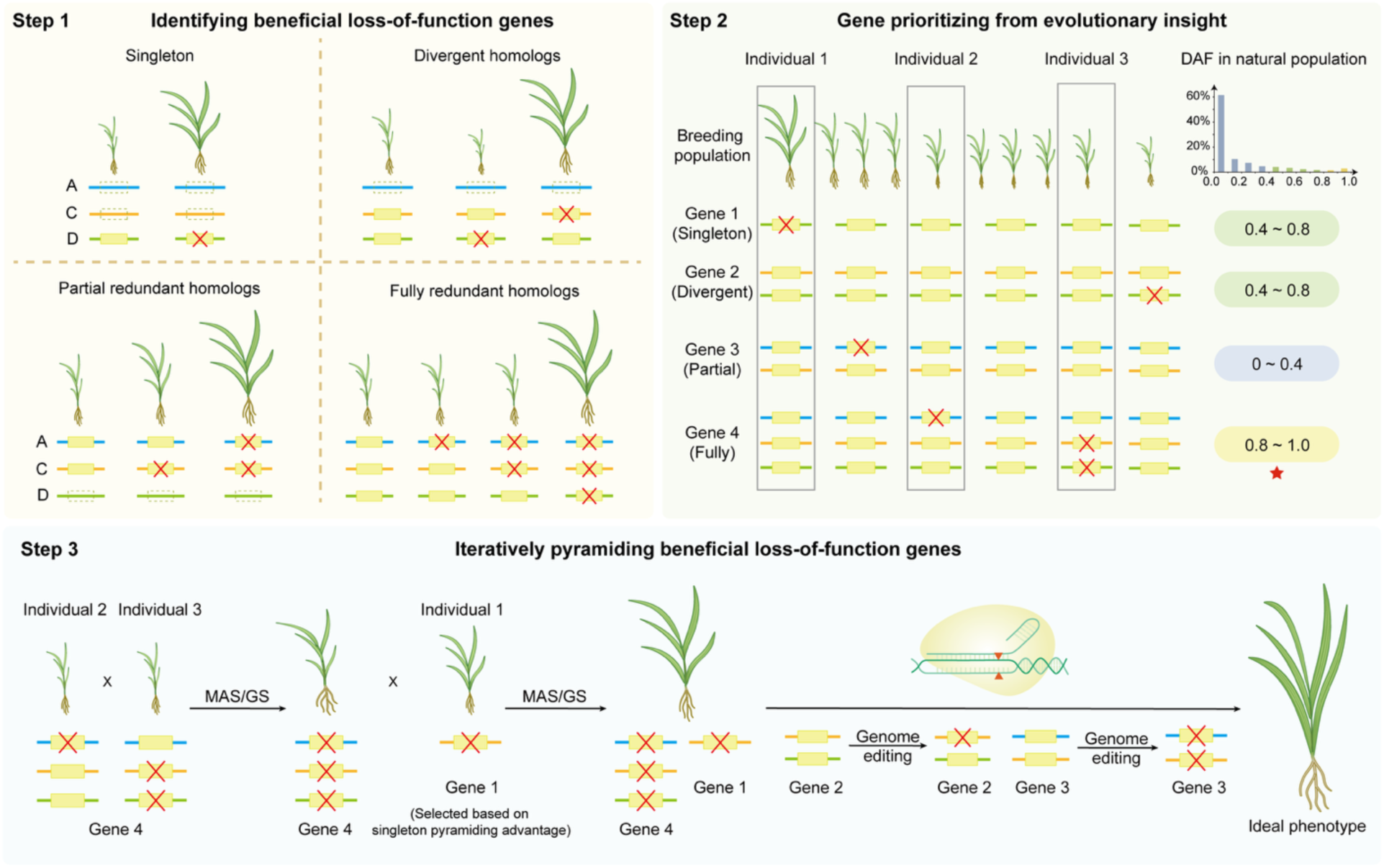
The targeted breeding roadmap for beneficial loss-of-function genes in hexaploid oat. To overcome the constraints posed by functional redundancy and buffering among homologs in hexaploid oat, we propose a breeding strategy that leverages natural variation and targeted genome editing. **Step 1**: Genes are classified into singletons, divergent homologs, partially redundant homologs, and fully redundant homologs through genomic screening. **Step 2**: Candidate genes are prioritized based on evolutionary insights, including derived allele frequency (DAF), nucleotide diversity, and evidence of convergence in natural populations. High-DAF deleterious alleles under positive selection are highlighted as prime targets for trait improvement. **Step 3**: Selected alleles are introduced and combined using marker-assisted selection (MAS), genomic selection (GS), and genome editing, with beneficial loss-of-function mutations pyramided through sequential fixation to ultimately achieve the desired phenotype.

For singletons, divergent homologs, and partially redundant homologs, which present relatively straightforward opportunities for direct trait modification, prioritizing the exploitation of natural variation appears advantageous. Marker-assisted selection (MAS)^33^ or genomic selection (GS)^34^ can be employed to enrich favorable alleles and accelerate genetic gain. In cases where natural variation appears limited, minimal genome editing approaches offer a precise means to introduce desirable modifications. By contrast, fully redundant homologs, where loss-of-function mutations are desirable, pose greater challenges due to the buffering effects of polyploidy. To address this, pyramiding beneficial loss-of-function alleles one triad at a time through sequential fixation with recurrent selection in manageable population sizes offers an effective strategy. When desirable loss-of-function alleles are absent, multiplex genome editing provides a powerful alternative to simultaneously disrupt all three homoeologous copies, thereby circumventing genetic redundancy with precise control. Such targeted genome editing increases the probability of achieving the intended phenotypic outcomes while minimizing unintended consequences.

## Discussion

The genus-level genome assemblies presented here offer new insights into longstanding questions regarding *Avena* evolution. Our analyses demonstrate that *A. agadiriana* is an allotetraploid comprising two distinct A subgenomes (A_g_ and A_gʹ_), which have experienced extensive inter-subgenomic chromosomal rearrangements. We also uncover evidence of hybrid speciation among diploid A genome species and observe substantial gene flow across species boundaries, revealing that the evolutionary history of *Avena* is more reticulate and dynamic than previously appreciated^12,20,35^. In contrast to patterns observed in some other polyploid crops^36–40^, hexaploid oat does not exhibit subgenome dominance. Instead, the A and D subgenomes show greater similarity in gene expression profiles and mutation loads than the more divergent C subgenome, and this subgenomic coherence contributes directly to the substantial genetic redundancy observed in the oat genome.

While such redundancy may help mask genetic burden and enhance genomic stability, it also presents challenges for breeding. The presence of multiple gene copies, coupled with subfunctionalization and dosage effects, can buffer both beneficial and deleterious mutations, thereby reducing the efficacy of selection. This buffering likely contributes to phenotypic plasticity in complex traits and may obscure the genetic signals needed to identify causal variants.

Although this study provides a foundational genomic framework, some questions remain. For example, the asymmetrical nucleotide diversity among the A, C, and D subgenomes in hexaploid oat may be the result of post-polyploidization introgression from diploid and tetraploid lineages— resembling patterns observed in wheat^41^—though further work is needed to clarify the timing, direction, and impact of such gene flow. Similarly, the extent to which genomic redundancy and subgenome interactions influence agronomic traits is not yet fully understood.

The proposed roadmap for targeted breeding highlights several conceptual and technical challenges that must be addressed. While genome editing and marker-assisted selection hold promise, their effectiveness depends on a detailed understanding of gene dosage, epistasis, and recombination landscapes—factors that remain under-characterized in oats. Moreover, the identification and prioritization of loss-of-function alleles that are both phenotypically advantageous and biologically tolerated are still in the early stages. We anticipate that the integration of high-resolution functional genomics, comparative pangenomics, and advanced phenotyping will be instrumental in decoding the complex genetic architecture of oats. While substantial challenges remain, we hope that our findings contribute meaningfully to ongoing efforts to better understand the evolutionary dynamics of polyploidy and to inform future research in *Avena* and other polyploid crops.

---

**Extended Fig. 1.**
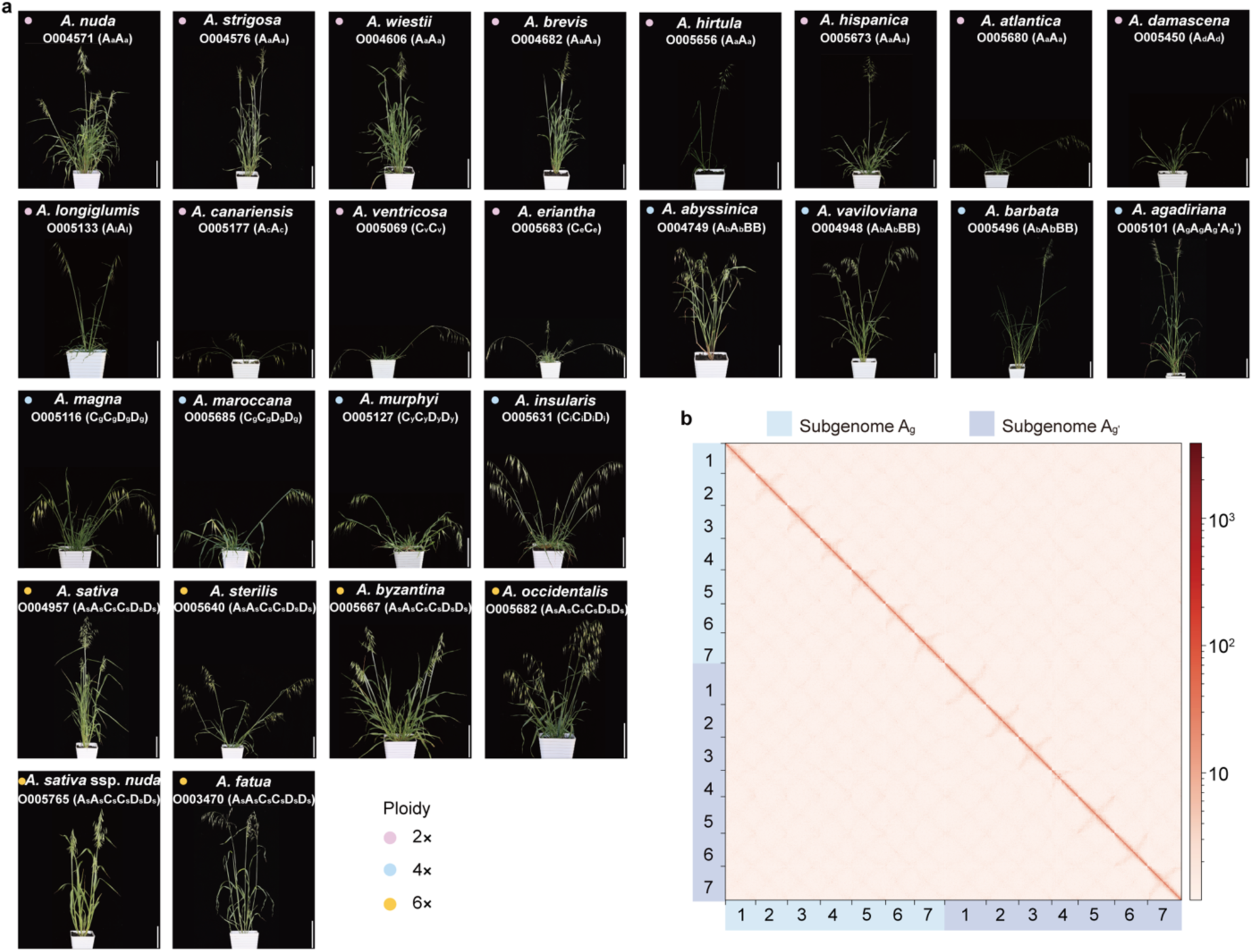
Overview of the diversity of the 26 *Avena* species and/or subspecies and Hi-C contact maps of *A. agadiriana*. **a**, Phenotypes of *Avena* species and/or subspecies for genome assembly. The information in parentheses indicate subgenome types. Scale bars = 18 cm. **b,** Hi-C contact maps of the 14 chromosomes of tetraploid *A. agadiriana*.

**Extended Fig. 2.**
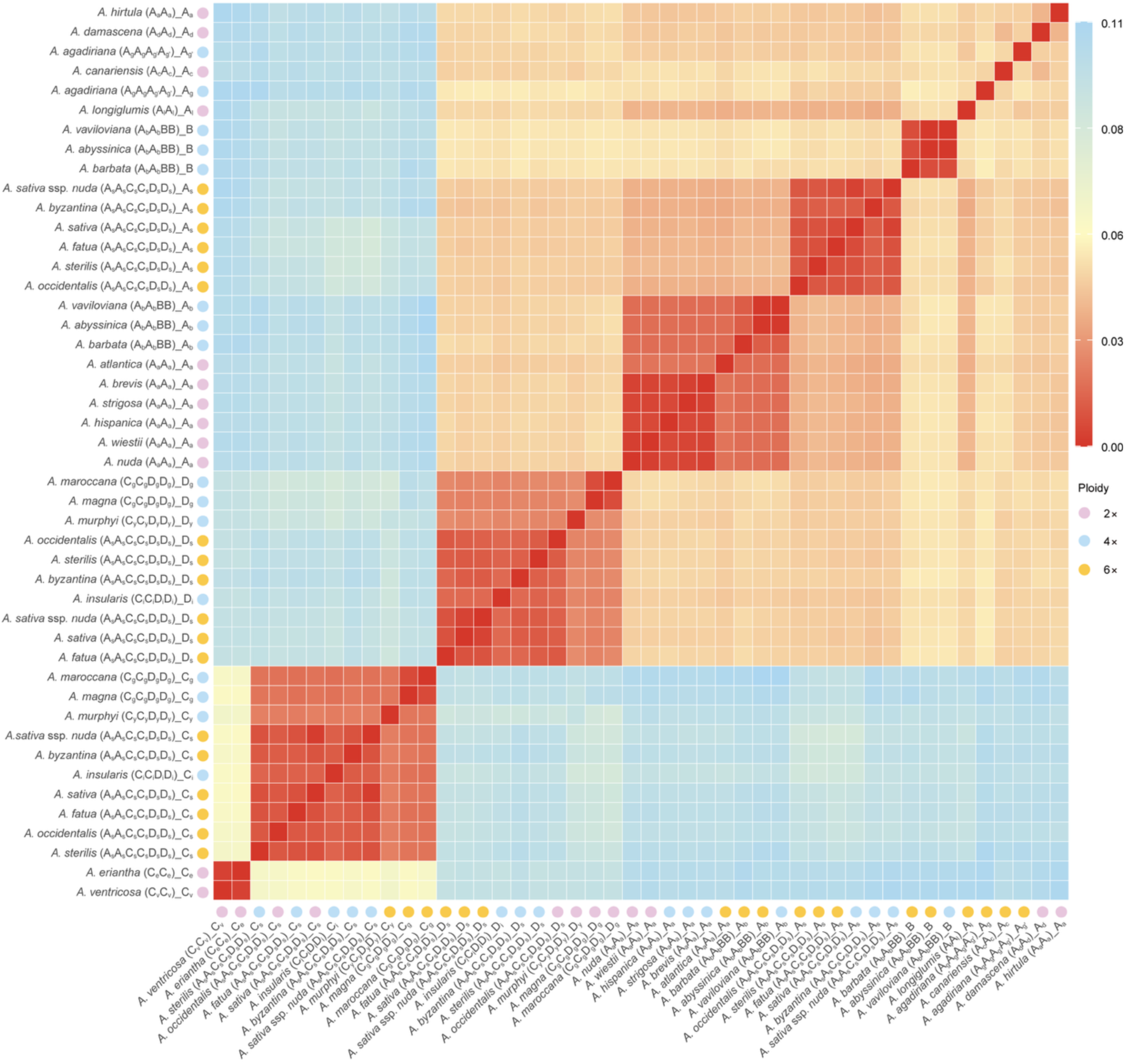
Pairwise genetic distance among the 46 subgenomes of the 26 *Avena* species and/or subspecies. Lower values represent closer genetic distances between subgenomes.

**Extended Fig. 3.**
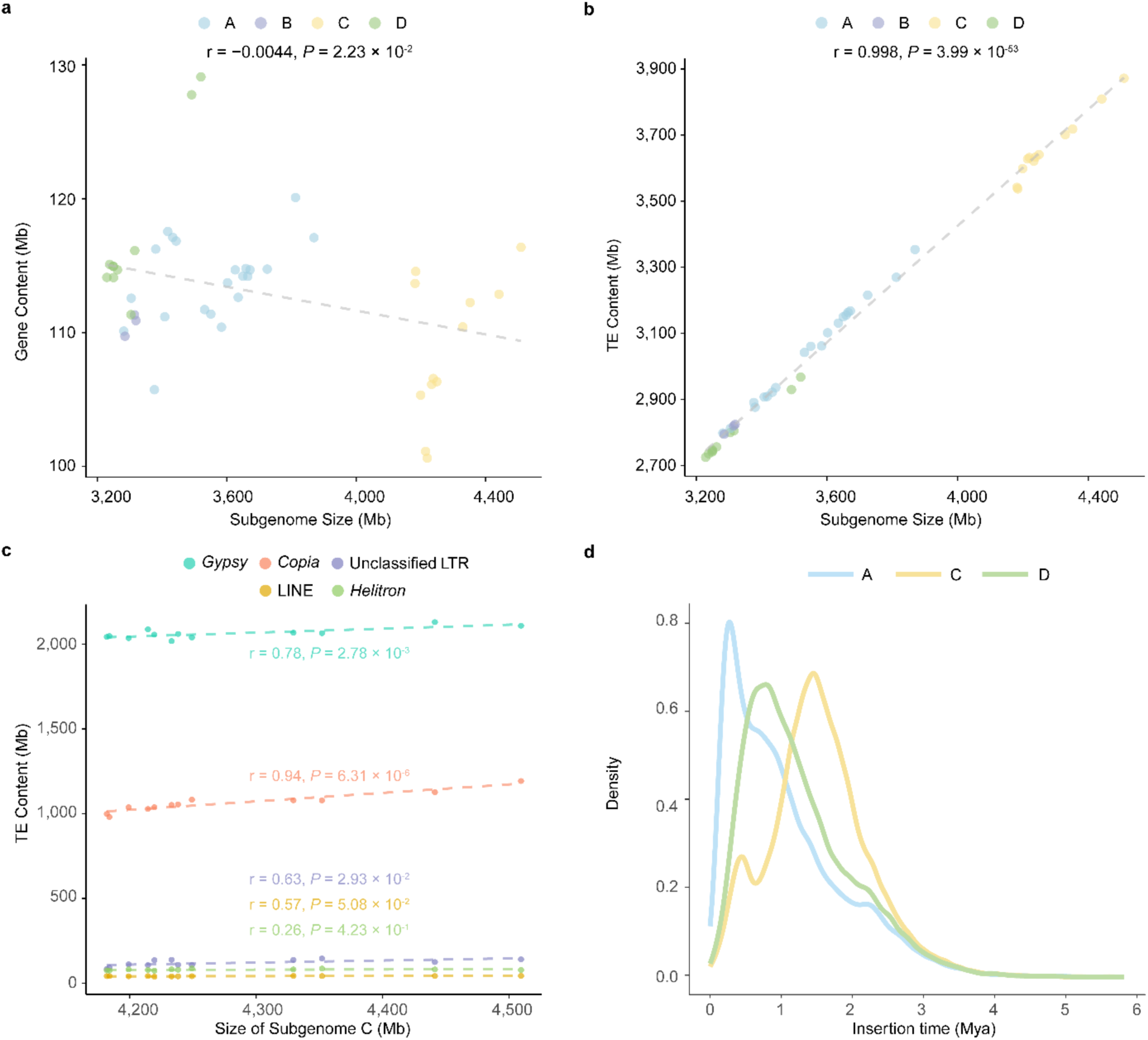
The correlations between subgenome size, gene content and transposable element (TE) content. **a, b**, Correlations between subgenome size and gene content (**a**), and between subgenome size and TE content (**b**). Each point represents a subgenome and is colored by subgenome type. **c,** Correlation between C subgenome size and the different content of class Ι TE family with individual family content exceeding 1%. Each point represents a type of TE and is colored by TE type. **a–c,** Pearson’s correlation coefficient was calculated, and its statistical significance was determined via a two-tailed *t*-test. **d,** Distribution of insertion times of TE in the hexaploid A, C, and D subgenomes. The peaks indicate periods of intensified transposon activity.

**Extended Fig. 4.**
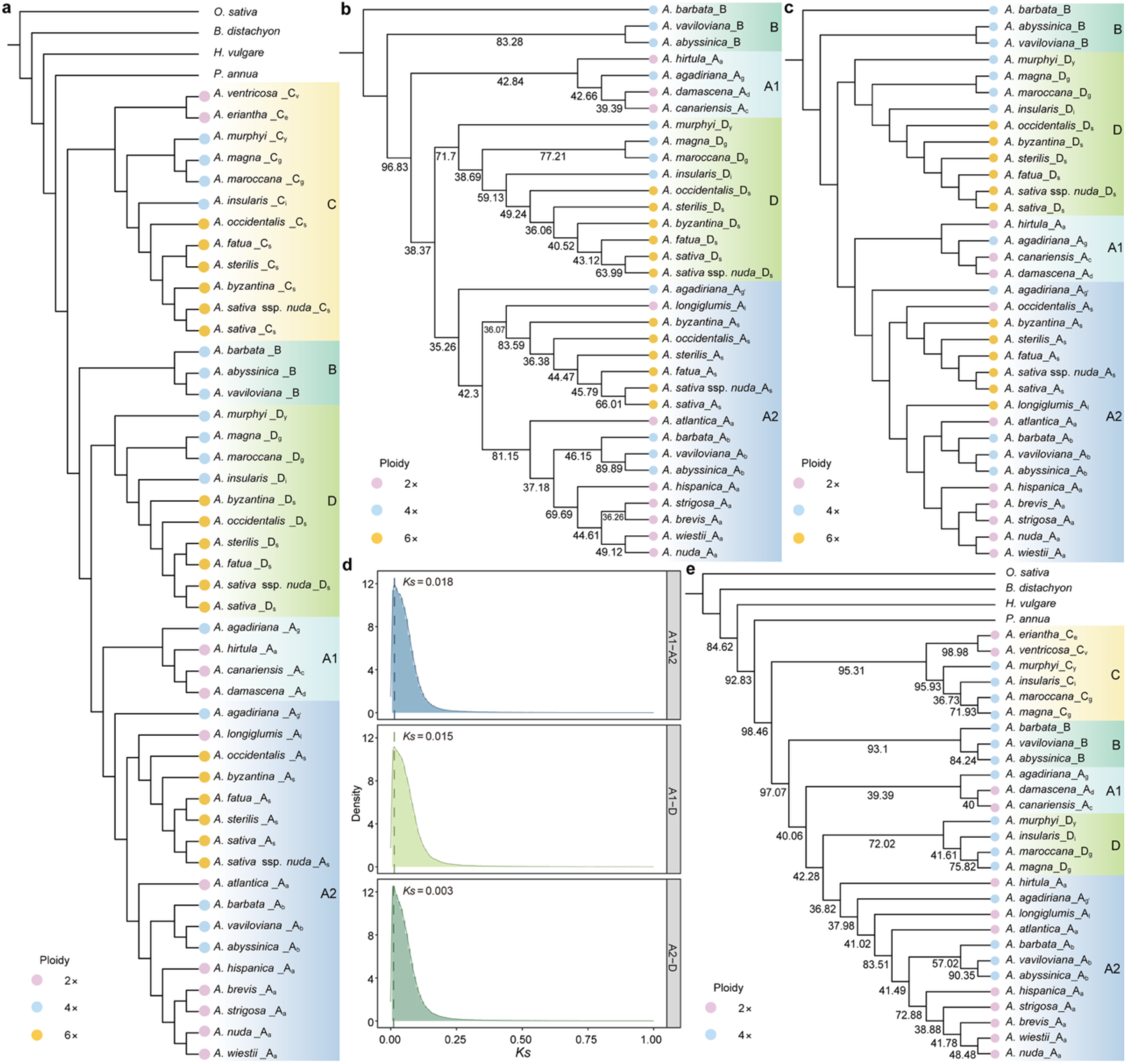
Phylogenetic analysis of *Avena* using different methods and datasets. **a**, Phylogenetic relationships among the 46 oat subgenomes and four Poaceae outgroups, reconstructed from 586 single-copy genes (SCGs) using the CASTER-pair^22^ method. The 46 oat subgenomes are grouped into five major clades (C, B, D, A1, and A2), each distinguished by different background colors. Branch lengths are not scaled to reflect evolutionary distances. **b, c,** Phylogenetic analyses using 2,889 SCGs identified across the A and D subgenomes, with the B subgenome as the outgroup, based on coalescent (**b**) and CASTER-pair^22^ (**c**) approaches. In panel **b**, the numbers on the nodes represent local posterior probability (localPP) values inferred from ASTRAL-III based on 2,889 individual gene trees. Branch lengths are not scaled to reflect evolutionary distances. **d,** Distribution of synonymous substitution rates (*Ks*) among A1–A2, A1–D, and A2–D clades. **e,** Phylogenetic reconstruction among the 28 oat subgenomes and four Poaceae outgroups, excluding hexaploidy individuals, based on the coalescent method using 586 SCGs. The numbers on the nodes represent local posterior probability (localPP) values inferred from ASTRAL-III based on 586 individual gene trees. Branch lengths are not scaled to reflect evolutionary distances.

**Extended Fig. 5.**
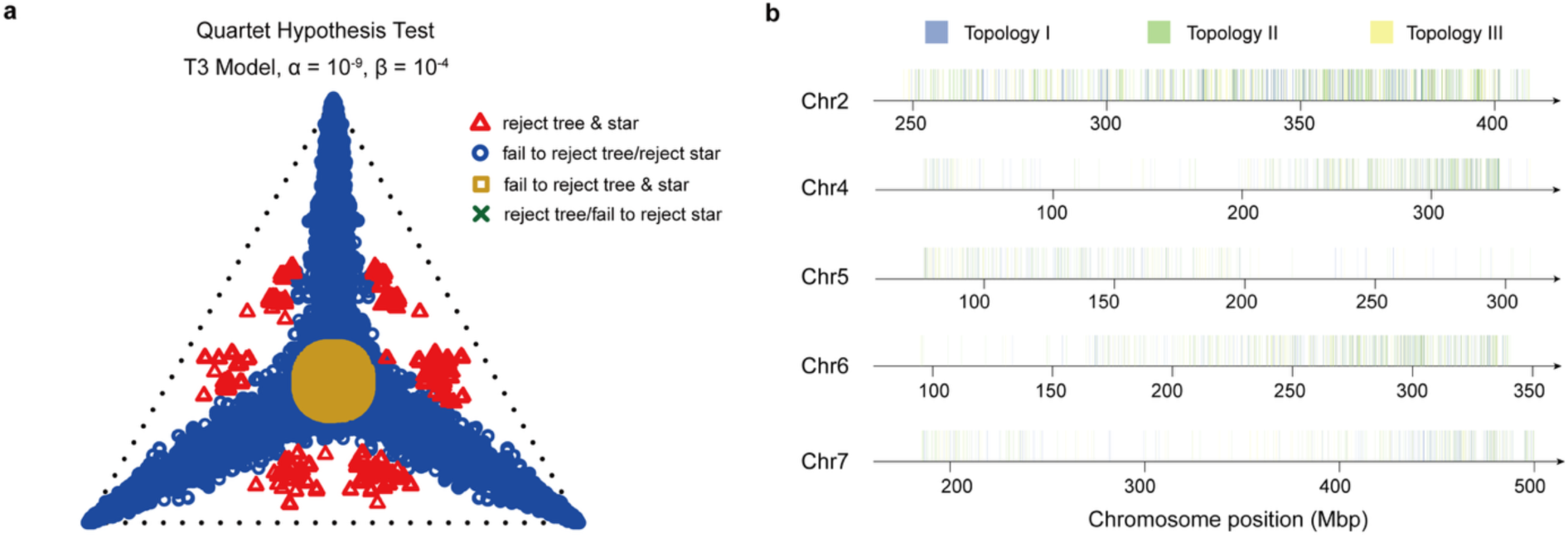
Test for ILS and introgression. **a**, Quartet Hypothesis Test under the T3 Model. The ternary plot represents the results of the quartet hypothesis test under the T3 model, with significance thresholds set at α = 1 × 10^-9^ and β = 1 × 10^-4^. Different symbols indicate the outcomes of hypothesis testing, indicating whether the null hypothesis is rejected or not. **b,** Genome-wide distribution of Topologies Ⅰ (purple bars), Ⅱ (green bars), and Ⅲ (yellow bars).

**Extended Fig. 6.**
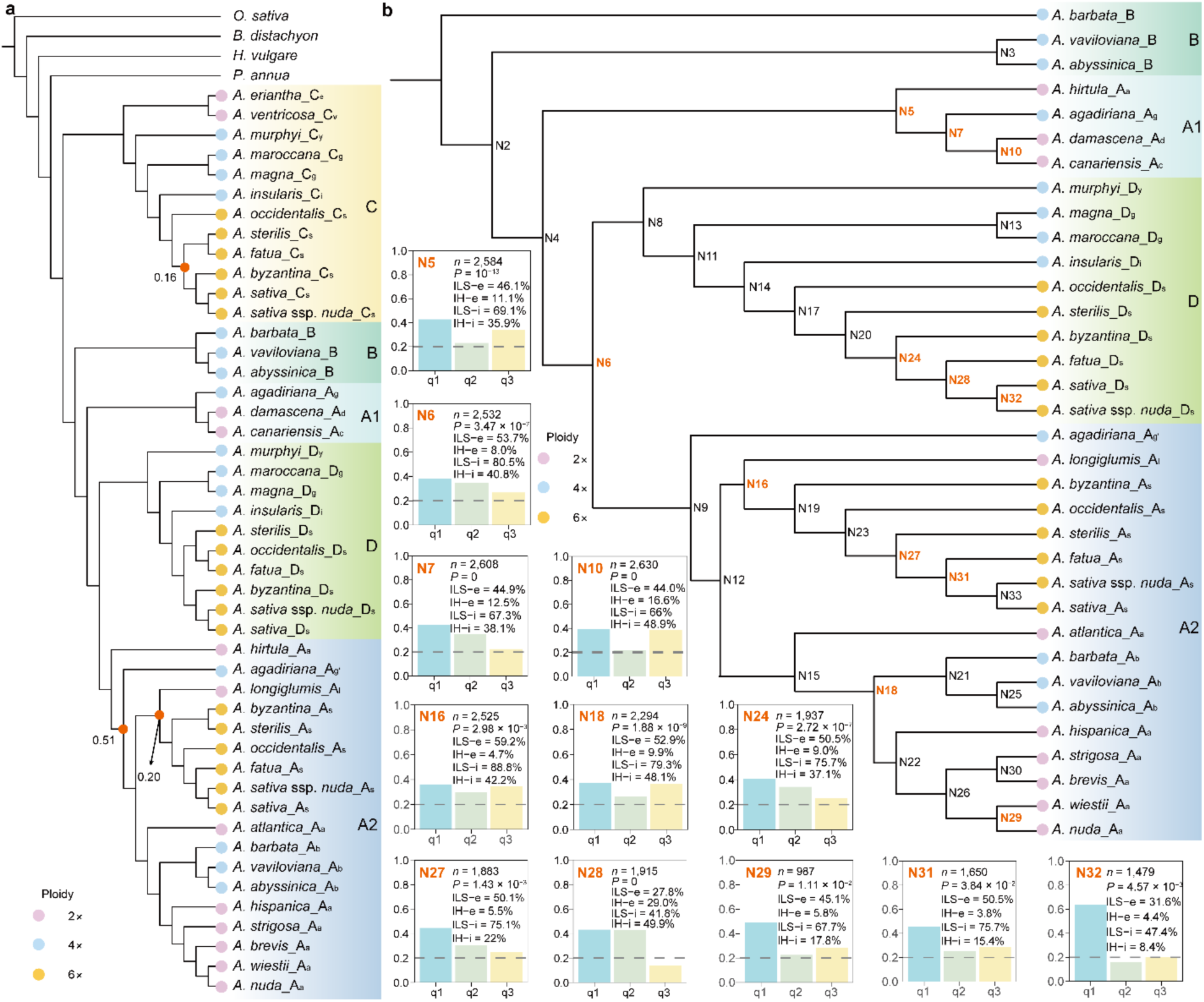
Extensive discordance between species tree and gene trees. **a**, Polytomy test based on the phylogenetic tree constructed from 586 single-copy genes (SCGs). *O. sativa*, *B. distachyon*, *H. vulgare*, and *P. annua* were used as outgroups. Shallow nodes with *P*-values greater than 0.05 are marked by red points, indicating that the null hypothesis of hard polytomy could not be rejected. **b,** Detection of incomplete lineage sorting (ILS) and introgression/hybridization (IH) indices at the nodes of 2,889 phylogenetic tree, with the B subgenome as the outgroup. N2–N33, labeled beside the nodes, represent node numbers. Nodes that cannot be solely attributed to ILS are marked in red. Detail detection results obtained by Phytop^42^ are shown to the left of the phylogenetic tree. *n* denotes the number of simulated gene trees, while *P* represents the *p*-value from the *χ^2^* test assessing whether the counts of topologies q2 and q3 are equal, ILS-e and IH-e indicate the proportions of gene tree topological incongruence attributable to ILS and IH, respectively. ILS-i and IH-i correspond to the calculated indices for ILS and IH.

**Extended Fig. 7.**
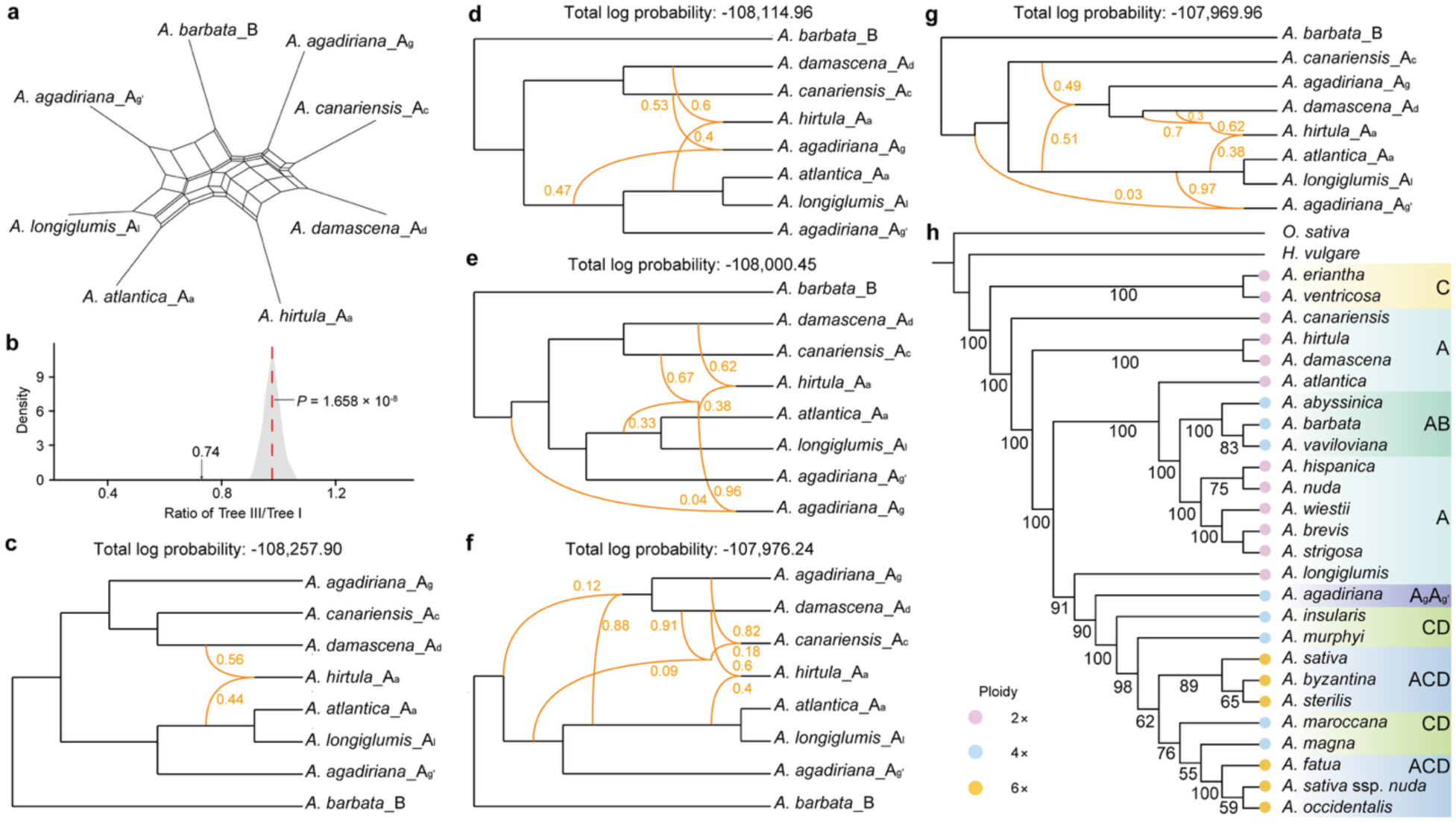
Reticulate evolution among A-genome diploids and chloroplast phylogenetic tree. **a**, Phylogenetic network analysis revealed the reticulate evolutionary history of A-genome diploids. **b,** Comparison of the observed topology ratio with that from a simulated ILS-only scenario. The black arrow indicates the observed ratios (Tree III/Tree II), while the gray curve represents the ratio under the ILS-only scenario. The ILS-only hypothesis was rejected based on two-tailed Student’s *t*-test. **c–g,** The hybridization scenarios of A-genome diploids inferred from PhyloNet^25^ using a dataset of 2,889 genes. Hybridization edges are shown as orange lines, with numbers indicating the inheritance probabilities from each parent. The total log probability is displayed at the top of the figure. **h,** Phylogenetic analysis based on the whole chloroplast genome of 26 *Avena* species and/or subspecies. Numbers on the nodes indicate bootstrap (BS) values. Branch lengths are not scaled to reflect evolutionary distances.

**Extended Fig. 8.**
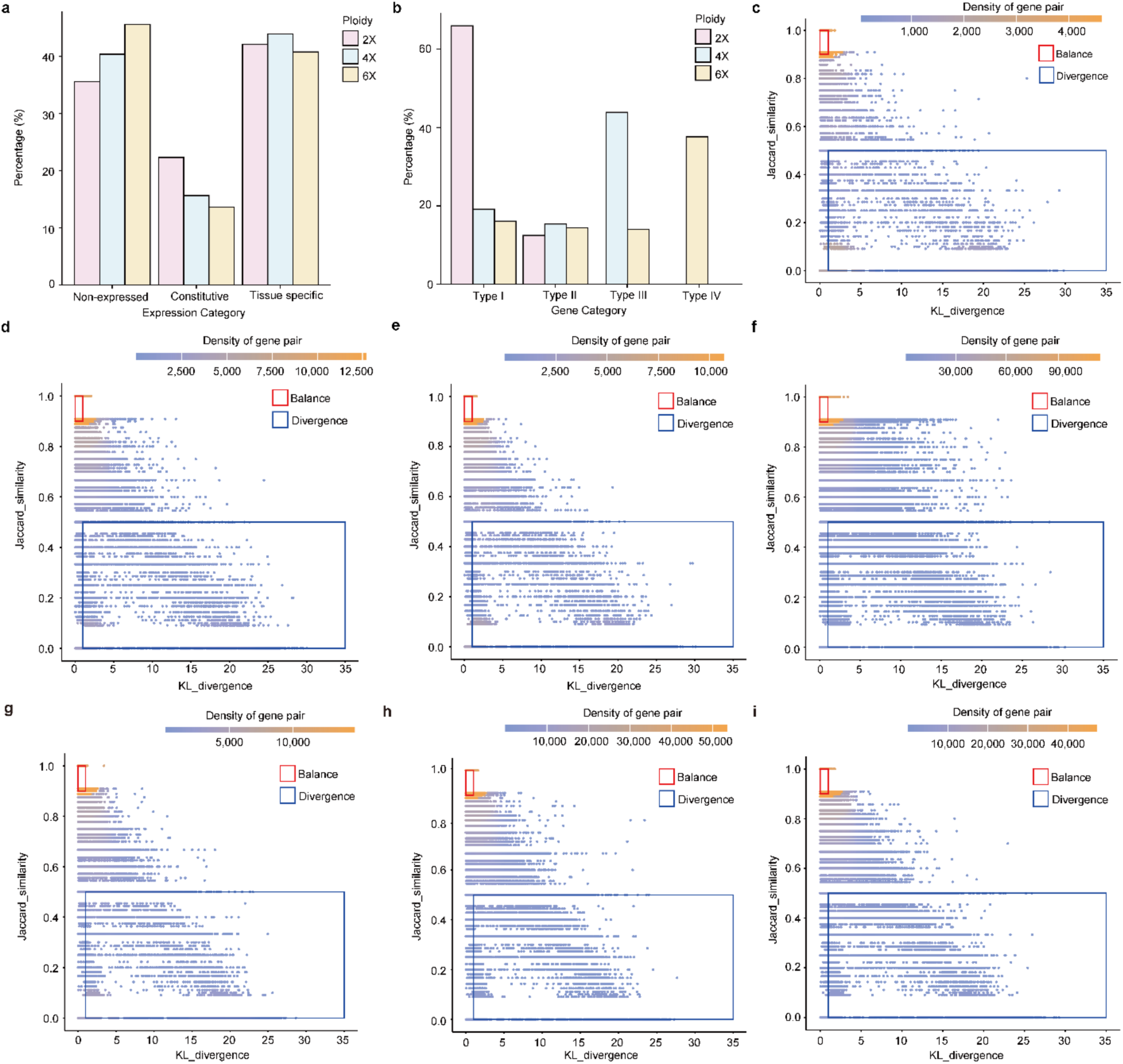
Gene expression pattern analysis across diploid, tetraploid and hexaploid oats. **a**, Proportions of Non-expressed, Constitutive, and Tissue-specific genes across diploid, tetraploid and hexaploid oats. **b,** Proportions of gene Types I–IV across diploid, tetraploid and hexaploid oats. **c–i,** Distribution of homologs based on their Kullback-Leibler (KL) divergence and Jaccard similarity in expression of paralogous pair in diploids (**c**), paralogous pair in tetraploids (**d**), paralogous pair in hexaploids (**e**), homoeologous pair in tetraploids (**f**), homoeologous pair in hexaploids (**g**), homoeologous traid (A–D subgenomes) of hexaploids (**h**), and homoeologous traid (C–D subgenomes) of hexaploids (**i**). Two expression groups were identified: Balance, defined by KL divergence in the range [0, 1] and Jaccard similarity in the range [0.9, 1.0]; and Divergence, defined by KL divergence in the range [1, 35] and Jaccard similarity in the range [0, 0.5].

**Extended Fig. 9.**
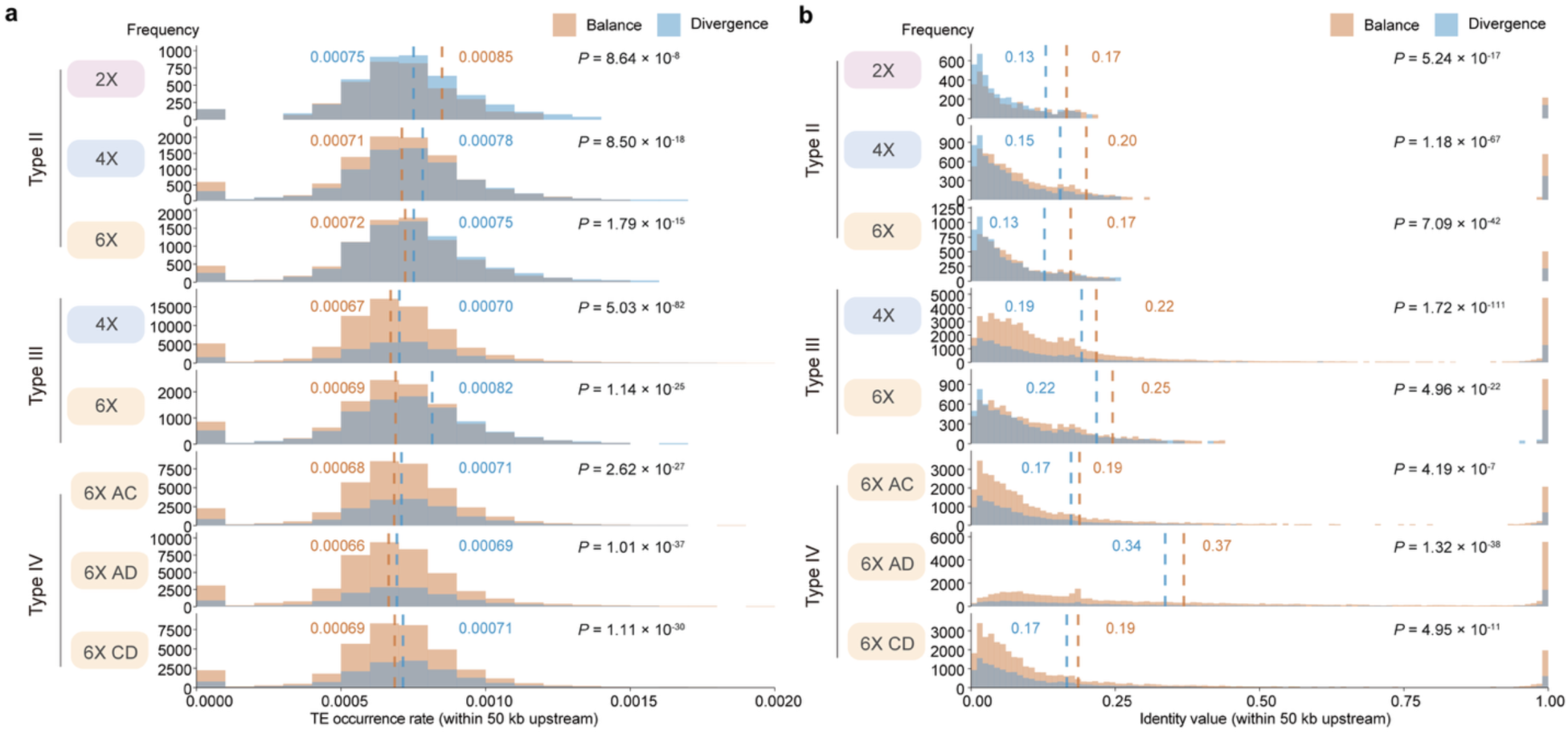
TE occurrence rate and sequence identity in 5’ upstream regions of balanced and divergent genes in Types II–IV. **a, b**, TE occurrence rate (**a**) and sequence identity (**b**) within the 50-kb 5’ upstream regions of balanced and divergent genes in Types II–IV. Dashed lines indicate mean values. *P*-values were calculated using Wilcoxon test.

**Extended Fig. 10.**
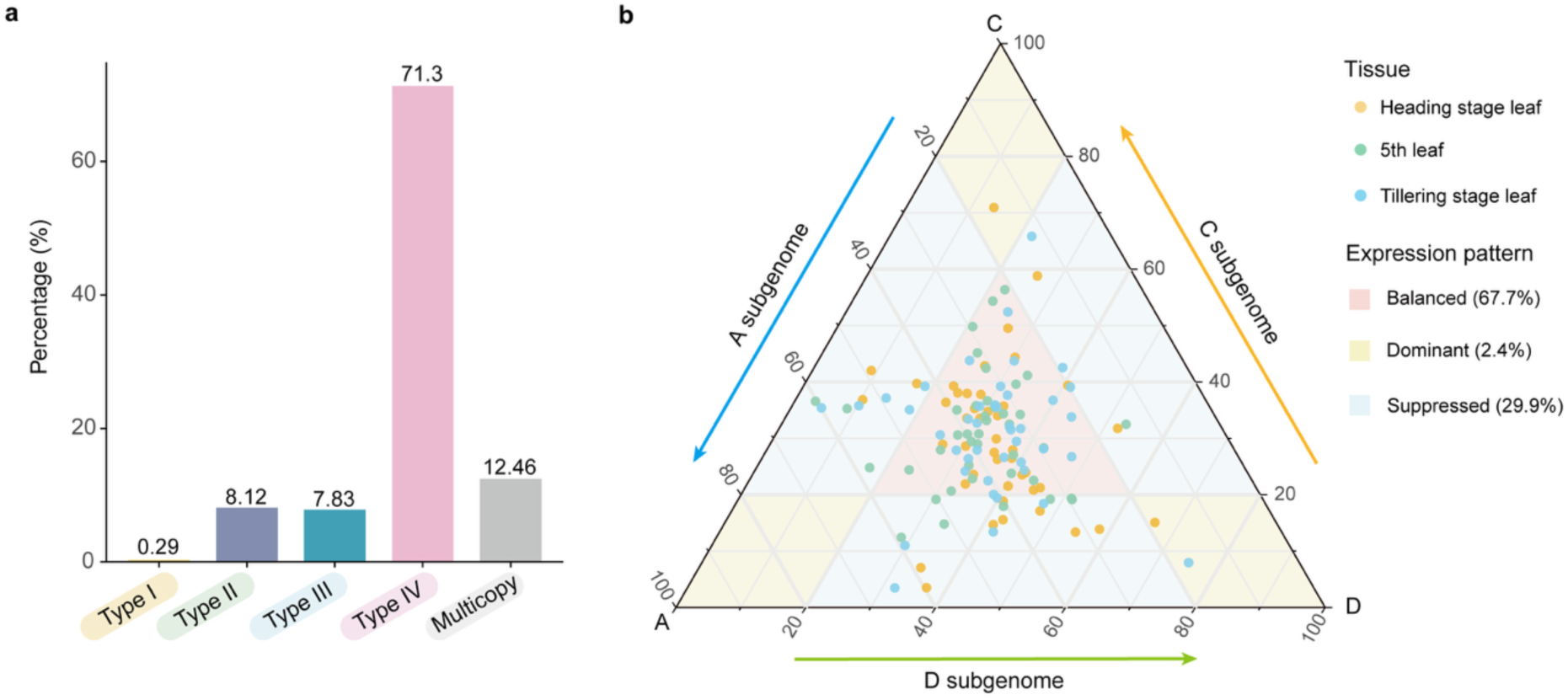
Identification and expression bias of leaf development gene orthologs in *A. sativa* ssp. *nuda*. **a**, Proportions of different gene types among the orthologs of 75 genes known to regulate leaf development in Arabidopsis, rice, wheat, and barley, identified in *A. sativa* ssp. *nuda*. **b,** Ternary plot of homoeologous triads for expression bias.

## Methods

### DNA preparation, sequencing and genome assembly

Genomic DNA from 26 *Avena* species and/or subspecies was extracted from fresh leaves. SMRTbell libraries were constructed following the manufacturer’s instructions from Pacific Biosciences and sequenced on the PacBio Revio sequencing platform. A total of 82 to 300 Gb of HiFi reads were generated using the PacBio official software CCS (https://github.com/PacificBiosciences/ccs). Hifiasm (v0.19.8)^43^ was used to assemble the HiFi reads into draft contigs with parameters (-l0 -f39), and the small, duplicated contigs, as well as plastid genomes^44,45^, were eliminated. Hi-C was performed following the protocol described previously^46^ with minor modifications. The final DNA library was prepared using the *Trans*NGS DNA Library Prep Kit for Illumina (kit ID: KP201) according to the manufacturer’s instructions. Libraries were quantified, phosphorylated and cyclized, and sequenced on the MGI DNBSEQ-T7 platform (PE150). The Hi-C data were used to obtain chromosome-level assemblies. Syntenic regions were identified using GENESPACE^47^ with default parameters, and the remaining assemblies were ordered and oriented.

### Protein-coding gene annotation

Helixer (v0.3.3)^48^, which integrates deep neural networks and hidden Markov models, was used to generate primary gene models in GFF3 format (--lineage land_plant). Subsequently, Liftoff (v1.6.3)^49^ was employed to map the annotation files of Sang^11^ and OT3098 (https://wheat.pw.usda.gov/jb?data=/ggds/oat-ot3098v2-pepsico) onto these 26 genomes with default parameters, producing corresponding GFF3 annotation files for each genome. Finally, EVidenceModeler (v2.1.0)^50^ was utilized to integrate these three annotation files with parameters (--segmentSize 100000 --overlapSize 10000), resulting in a high-confidence genome annotation.

### Subgenome distance calculation, SubPhaser and identification of syntenic blocks

To estimate the genetic distance between subgenomes, the oat tetraploid and hexaploid genomes were partitioned. Mash (v2.1)^51^ was then employed to construct sketches for the 46 subgenomes using *mash sketch* and to calculate pairwise genetic distances with *mash dist*. The subgenome distances were subsequently visualized using the R pheatmap package. To further validate the subgenome assignments, SubPhaser (v1.2.6)^52^ (-disable_ltr -disable_circos) was utilized for subgenome-specific k-mer detection. The identified k-mers were then used for principal component analysis (PCA) and clustering heatmap visualization using R package. To identify syntenic blocks across all subgenomes of 26 *Avena* species and/or subspecies, we employed the WGDI pipeline (v0.6.5)^53^ to detect complete homoeologous blocks. Subsequently, we used BLASTP (v2.12.0)^54^ with default parameters to compute pairwise sequence similarities, using the A subgenome of *A*. *sativa* ssp. *nuda* as the reference.

### Phylogenetic analyses

To construct the subgenome phylogenetic tree of 26 *Avena* species and/or subspecies, amino-acid sequences were initially extracted from the subgenomes of 26 *Avena* and 4 outgroups (*O*. *sativa*, *B*. *distachyon*, *H*. *vulgare*, and *P*. *annua*) using gffread (v0.12.7)^55^ with parameters (-y). Subsequently, the resultant sequences results were processed through Sonicparanoid2 (v2.0.8)^56^ with default parameters to infer orthology. A total of 586 single-copy orthologous genes within each subgenome, all residing in syntenic blocks, were obtained. Then, protein alignments for these genes were computed using MAFFT (v7.525)^57^ with parameters (mafft-linsi --maxiterate 1000 -- localpair). The protein alignments were then converted to their corresponding coding sequences (CDS), where each amino acid was replaced by its respective triplet bases from the CDS based on identical ID information, employing PAL2NAL2 (v14)^58^ with default parameters. The resulting locus alignment files were refined using trimAl (v1.4.rev15)^59^ with parameters (-gt 0.75). For each locus, maximum likelihood trees were generating using IQ-TREE (v1.6.9)^60^. Node support was established through 1,000 ultrafast bootstrap (UFBS) replicates, considering UFBS values ≥ 95% as indicative of strong support. iTOL (<u>iTOL: Interactive Tree of Life (embl.de)</u>^61^ was used to visualize the result of IQ-TREE (v1.6.9) with parameters (-m GTR-bb 1000 -bnni -redo -nt AUTO -pre). To minimize the effect of ILS, a multi-species coalescent-based method implemented in ASTRAL-III (v5.7.8)^62^ was applied to infer the species tree. Edges of the input gene trees with UFBS ≤ 30 collapsed using Newick Utilities (v1.6.0)^63^ with parameters (’i & b<=30’). Branch support was assessed using localPP^64^, accessed by ASTRAL-III (v5.7.8)^62^ with parameters (-t1). Additionally, the CASTER-pair^22^ method was also employed to reconstruct the species tree with default parameters. Divergence time was estimated using MCMCTREE algorithm from PAML package (v4.5)^65^ based on the 586 SCGs phylogenetic relationships inferred by ASTRAL-III^62^. One calibration time from the TimeTree database (http://www.timetree.org/) for the divergence between *O*. *sativa* and *B*. *distachyon* (41.5–62.0 Mya) and another calibration time from a previous study^66^ for the divergence between *B*. *distachyon* and *H*. *vulgare* (39.9–40.4 Mya), *H*. *vulgare*, and *P*. *annua* (35.5–35.9 Mya), *P*. *annua* and *A*. *sativa* (28.7–29.1 Mya) were used for molecular dating.

### Conflict calculation between gene trees and species trees

To systematically evaluate phylogenetic discordance between species tree and individual gene trees, we employed MSCquartets^67^ to analyze conflicts between species trees and gene trees. Initially, we calculated the quartet count concordance factor (qcCF) for each set of four taxa across all gene trees, identifying the frequencies of the three possible topologies. By comparing these frequencies, *P*-values were derived to evaluate the congruence between gene trees and the species tree. The qcCF for all quartets was visualized using simplex plots under the T3 model, which does not require a predefined species tree. Using 586 SCGs trees, the plots were generated. Using the 586 SCGs tree as a reference, the conflicts between the individual gene trees and the species tree were calculated by phyparts^68^ with default parameters. Only branches in the gene trees with UFBS values greater than 30% were displayed. The results were visualized using the Python script ‘phypartspiecharts.py’ (<u>phyloscripts/phypartspiecharts at master mossmatters/phyloscripts</u> <u>GitHub</u>). Based on the results from ASTRAL-III (v5.7.8)^62^ with parameters (-t2), we calculated the ILS and hybridization signals at each node using Phytop^42^ with default parameters. Additionally, we employed twisst^69^ to statistically analyze the topological structures among different taxa with default parameters.

### Gene flow and Polytomy test

We employed the tree-based method QuIBL (https://github.com/miriammiyagi/QuIBL)^24^ to distinguish introgression from ILS. The input files from the fasta alignments of 2,889 SCGs were generated using IQ-TREE (v1.6.9)^60^ with parameters (-m GTR -bb 1000 -bnni -redo -nt AUTO - pre). We then compared an ILS-only model to an introgression + ILS model using the Bayesian Information Criterion (BIC), with a strict cutoff of ΔBIC > 10. The QuIBL results were subsequently analyzed using the quiblR package (https://github.com/nbedelman/quiblR). To access whether a node in the species tree should be represented as a polytomy, the polytomy test^70^ integrated into ASTRAL-III (v5.7.8)^62^ were performed with parameters (-t 10).

### Phylogenetic network analyses

To access whether reticulate evolution contributes to the observed conflicts, we employed the Network inference Algorithm via NeighbourNet Using the Quartet distance (NANUQ)^71^ method to generate a network. Using the MSCquartets R package^67^, We calculated empirical quartet counts from the gene trees and determined network quartet distances between taxa with parameters (“α = 0.01” and “β = 0.95”) to construct a likelihood network. Additionally, we utilized the maximum pseudo-likelihood method in PhyloNet (v.3.8.2)^25^ to construct a phylogenetic network based on the 2,889 SCGs representing the 3-taxon data, using the parameters (InferNetwork_MPL (all) -d). Our network searches allowed for 1–5 reticulations and included only nodes from the rooted maximum likelihood (ML) gene trees with ultrafast bootstrap (UFBoot) support of at least 30%. The networks were evaluated based on their total log probability.

### ILS simulation

Simulation of 2,889 gene trees with ILS among 26 *Avena* species and/or subspecies were performed by DendroPy (v4.2.0)^72^. The species tree, inferred using ASTRAL-III^62^ from 2,889 SCGs, served as the input. The frequencies of observed gene-tree topologies with those from the simulations were compared to evaluate if ILS alone could explain the phylogenetic discordance.

### Transcriptome quantification, homolog classification and expression pattern analysis of homoeologous triads

For each of the 26 species and/or subspecies, transcriptome data were generated from 11 distinct tissues. A total of 286 raw RNA-seq datasets were quality-filtered using fastp (v0.24.0)^73^ to remove adapter sequences, low-quality bases, and duplicated reads with parameter (--dedup). The quality of the filtered reads was subsequently assessed using FastQC (http://www.bioinformatics.babraham.ac.uk/projects/fastqc/, v0.12.1) and MultiQC (v1.25.2)^74^. To ensure sufficient and reliable data coverage, filtered minimum clean data amounts were established according to ploidy level: 3 Gb for diploids, 6 Gb for tetraploids, and 9 Gb for hexaploids. Transcript quantification was performed using kallisto (v0.51.1)^75^ with default parameters, using the genome assembly of each respective species as the reference for index construction. Gene expression levels were measured in transcripts per million (TPM), and genes with TPM > 1 were considered actively expressed in each sample.

To classify homologous gene groups in *Avena*, we integrated two complementary reference gene sets. The first was generated using SonicParanoid2 (v2.0.8)^56^ with default parameters, which identified orthogroups from coding sequence (CDS) files across 46 subgenomes in the genus. The second gene set was produced using FastOMA (v0.3.4)^76^ with default parameters. Gene groups were categorized into four main categories based on gene copy number and their distribution across the A, C, and D subgenomes: (1) Singleton: a single gene copy present in any one subgenome. (2) Paralogous pair: two gene copies present within the same subgenome. (3) Homoeologous pair: one gene copy in each of two subgenomes. (4) Homoeologous triad: one gene copy in each of the A, C, and D subgenomes. Gene groups that did not fit any of these categories were classified as multi-copy types. Classification was based on the integration of both datasets: paralogous pairs were inferred from SonicParanoid2 (v2.0.8)^56^, while singletons, homoeologous pairs, and homoeologous triads were defined primarily using FastOMA (v0.3.4)^76^, with supplementary support from SonicParanoid2 (v2.0.8)^56^.

To characterize the expression patterns of homoeologous triads related to leaf development, we focused on orthologs previously reported to regulate leaf development in Arabidopsis, rice, wheat, and barley. We analyzed their expression in leaf tissues collected at the five-leaf, tillering, and heading stages. Homoeologous triads were extracted from the constructed expression matrix, and only these with a total expression (TPM-A + TPM-C + TPM-D) greater than 1 were retained for analysis. To assess the relative contribution of each subgenome within these triplets, we calculated the normalized expression proportion using the following formula (where A, C, and D denotes the gene IDs of the homoeologs in subgenomes A, C, and D):

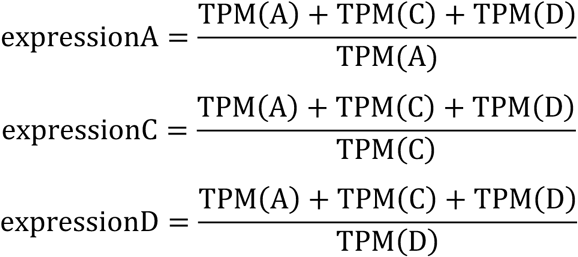

### Identification of deleterious mutation

To identify deleterious mutations, whole-genome resequencing data from oat samples^30^ were first aligned to the reference genome *A. sativa* ssp. *nuda* using BWA (v0.7.17)^77^ with default parameters. The resulting alignments were sorted and processed using DeepVariant (v1.6.1)^78^ with default parameters to perform single-sample variant calling, yielding a VCF file for each accession. Subsequently, these individual VCF files were then jointly genotyped and merged using GLnexus (v1.1.1)^79^, generating a combined VCF file containing all variant sites across the sampled accessions after filtering out low-confidence variants. To evaluate the functional impact of the variants, the filtered VCF file—restricted to cultivated accessions—was annotated using SIFT 4G^31^. A custom SIFT 4G annotation database was built for the *A. sativa* ssp. *nuda* reference genome using the angiosperm dataset from UniProt90 (https://www.uniprot.org/). SIFT was run with default parameters to predict functional consequences and assign SIFT scores. Variants with SIFT scores < 0.05 were classified as deleterious and retained for downstream functional impact and statistical analyses.

## Reporting summary

Further information on research design is available in the Nature Portfolio Reporting Summary linked to this article.

## Data and code availability

All sequence data of this study have been deposited at the Sequence Read Archive (https://ncbi.nlm.nih.gov/sra) under BioProject PRJNA1259686.

## Acknowledgements

We thank United States Department of Agriculture, Agriculture and Agri-Food Canada, Leibniz Institute of Plant Genetics and Crop Plant Research, and John Innes Centre for providing *Avena* germplasms. This work was supported by the Strategic Priority Research Program of the Chinese Academy of Sciences (XDA26030102), and the National Natural Science Foundation of China (32441029).

## Author contributions

Y. Zhou conceived and designed the research. X.B., Y. Zhang, X.C., X.L., D.W., Z.Z., J.H. and M.L. participated in material preparation. X.B. performed the Hi-C assay. H.Y. contributed to the genome assembly and gene annotation. M.S. contributed to the transposable element annotation. Y. Zhang contributed to the phylogenetic analysis, network analysis, and divergence time estimation. X.C. contributed to the identification of homoeologous exchange. Y. Zhou, M.S. and H.Y. contributed to the transcriptome analysis. M.S. contributed to the figure preparation. X.B. wrote the draft manuscript. Y. Zhou wrote and revised the manuscript. Y. Zhou supervised the research. Z.B., Z.Z., Z.D., Z.L., Y.W., L.Z. and L.Y. revised the final manuscript. All the authors read, edited and approved the manuscript.

## Additional information

Supplementary Information is available for this paper.

Correspondence and requests for materials should be addressed to Yao Zhou.

## References

1. Soltis, D. E., Soltis, P. S. & Tate, J. A. Advances in the study of polyploidy since plant speciation. New Phytol. 161, 173–191 (2004).

2. Sattler, M. C., Carvalho, C. R. & Clarindo, W. R. The polyploidy and its key role in plant breeding. Planta. 243, 281–296 (2016).

3. Salman-Minkov, A., Sabath, N. & Mayrose, I. Whole-genome duplication as a key factor in crop domestication. Nat. Plants. 2, 16115 (2016).

4. Renny-Byfield, S. & Wendel, J. F. Doubling down on genomes: polyploidy and crop plants. Am. J. Bot. 101, 1711–1725 (2014).

5. Fang, Z. & Morrell, P. L. Domestication: polyploidy boosts domestication. Nat. Plants 2, 16116 (2016).

6. Schiessl, S. V., Katche, E., Ihien, E., Chawla, H. S. & Mason, A. S. The role of genomic structural variation in the genetic improvement of polyploid crops. Crop J. 7, 127–140 (2019).

7. Van de Peer, Y., Mizrachi, E. & Marchal, K. The evolutionary significance of polyploidy. Nat. Rev. Genet. 18, 411–424 (2017).

8. Doyle, J. J. et al. Evolutionary genetics of genome merger and doubling in plants. Annu. Rev. Genet. 42, 443–461 (2008).

9. Van de Peer, Y., Maere, S. & Meyer, A. The evolutionary significance of ancient genome duplications. Nat. Rev. Genet. 10, 725–732 (2009).

10. Iohannes, S. D. & Jackson, D. Tackling redundancy: genetic mechanisms underlying paralog compensation in plants. New Phytol. 240, 1381–1389 (2023).

11. Kamal, N. et al. The mosaic oat genome gives insights into a uniquely healthy cereal crop. Nature 606, 113–119 (2022).

12. Peng, Y. et al. Reference genome assemblies reveal the origin and evolution of allohexaploid oat. Nat. Genet. 54, 1248–1258 (2022).

13. Baum, B. R. Oats: wild and cultivated. A monograph of the genus Avena L. (Poaceae). (Minister of Supply and Services, Ottawa, 1977).

14. Thomas, H. Cytogenetics of *Avena*. In: oat science and technology (eds Marshall, H. G. & Sorrells, M. E.) 473–507 (American Society of Agronomy, Crop Science Society of America, Madison, 1992).

15. Ladizinsky, G. Studies in oat evolution: a man’s life with Avena. Springer, Heidelberg (2012).

16. Jellen, E. N., et al. A uniform gene and chromosome nomenclature system for oat (*Avena* spp.). Crop Pasture Sci. 75, CP23247 (2024).

17. Yan, H. et al. High-density marker profiling confirms ancestral genomes of *Avena* species and identifies D-genome chromosomes of hexaploid oat. Theor. Appl. Genet. 129, 2133–2149 (2016).

18. Loskutov, I. G. On evolutionary pathways of *Avena* species. Genet. Resour. Crop Evol. 55, 211–220 (2008).

19. Chew, P. et al. A study on the genetic relationships of *Avena* taxa and the origins of hexaploid oat. Theor. Appl. Genet. 129, 1405–1415 (2016).

20. Tomaszewska, P., Schwarzacher, T. & Heslop-Harrison, J. S. P. Oat chromosome and genome evolution defined by widespread terminal intergenomic translocations in polyploids. Front. Plant. Sci. 13, 1026364 (2022).

21. Leggett, J. M. et al. Intergenomic translocations and the genomic composition of *Avena maroccana* Gdgr. revealed by FISH. Chromosome Res. 2, 163–164 (1994).

22. Zhang, C., Nielsen, R. & Mirarab, S. CASTER: direct species tree inference from whole-genome alignments. Science 387, eadk9688 (2025).

23. Wang, Z. et al. Genomic evidence for homoploid hybrid speciation between ancestors of two different genera. Nat. Commun. 13, 1987 (2022).

24. Edelman, N. B. et al. Genomic architecture and introgression shape a butterfly radiation. Science 366, 594–599 (2019).

25. Than, C., Ruths, D. & Nakhleh, L. PhyloNet: a software package for analyzing and reconstructing reticulate evolutionary relationships. BMC Bioinformatics 9, 322 (2008).

26. Adams, K. L. & Wendel, J. F. Novel patterns of gene expression in polyploid plants. Trends Genet. 21, 539–543 (2005).

27. Cheng, F. et al. Gene retention, fractionation and subgenome differences in polyploid plants. Nat. Plants 4, 258–268 (2018).

28. The GTEx Consortium et al. The genotype-tissue expression (GTEx) pilot analysis: multitissue gene regulation in humans. Science 348, 648–660 (2015).

29. Simons, C., Pheasant, M., Makunin, I. V. & Mattick, J. S. Transposon-free regions in mammalian genomes. Genome Res. 16, 164–172 (2006).

30. Nan, J. et al. Genome resequencing reveals independent domestication and breeding improvement of naked oat. Gigascience 12, giad061 (2022).

31. Vaser, R., Adusumalli, S., Leng, S. N., Sikic, M. & Ng, P. C. SIFT missense predictions for genomes. Nat. Protoc. 11, 1–9 (2016).

32. Aluko, O. O., Li, C., Wang, Q. & Liu, H. Sucrose utilization for improved crop yields: a review article. Int. J. Mol. Sci. 22, 4704 (2021).

33. Hasan, N., Choudhary, S., Naaz, N., Sharma, N. & Laskar, R. A. Recent advancements in molecular marker-assisted selection and applications in plant breeding programmes. J. Genet. Eng. Biotechnol. 19, 128 (2021).

34. Alemu, A. et al. Genomic selection in plant breeding: key factors shaping two decades of progress. Mol. Plant 17, 552–578 (2024).

35. Fu, Y. B. Oat evolution revealed in the maternal lineages of 25 *Avena* species. Sci. Rep. 8, 4525 (2018).

36. Cheng, F. et al. Biased gene fractionation and dominant gene expression among the subgenomes of *Brassica rapa*. PLoS One 7, e36442 (2012).

37. Yoo, M. J., Szadkowski, E. & Wendel, J. F. Homoeolog expression bias and expression level dominance in allopolyploid cotton. Heredity 110, 171–180 (2013).

38. Edger, P. P. et al. Subgenome dominance in an interspecific hybrid, synthetic allopolyploid, and a 140-year-old naturally established neo-allopolyploid monkeyflower. Plant Cell 29, 2150–2167 (2017).

39. Lovell, J. T. et al. Genomic mechanisms of climate adaptation in polyploid bioenergy switchgrass. Nature 590, 438–444 (2021).

40. Ma, P. F. et al. Genome assemblies of 11 bamboo species highlight diversification induced by dynamic subgenome dominance. Nat. Genet. 56, 710–720 (2024).

41. Zhou, Y. et al. *Triticum* population sequencing provides insights into wheat adaptation. Nat. Genet. 52, 1412–1422 (2020).

## References

42. Shang, H. Y. et al. Phytop: a tool for visualizing and recognizing signals of incomplete lineage sorting and hybridization using species trees output from ASTRAL. Hortic. Res. 12, uhae330 (2024).

43. Cheng, H., Concepcion, G. T., Feng, X., Zhang, H. & Li, H. Haplotype-resolved *de novo* assembly using phased assembly graphs with hifiasm. Nat. Methods 18, 170–175 (2021).

44. Wang, H. et al. Characterization of the complete chloroplast genome of *Avena chinensis* (Poales: Poaceae). Mitochondrial DNA B. Resour. 6, 3137–3139 (2021).

45. Liu, Q. et al. The mitochondrial genome of the diploid oat *Avena longiglumis*. BMC Plant Biol. 23, 218 (2023).

46. Huang, J., Jiang, Y., Zheng, H. & Ji, X. BAT Hi-C maps global chromatin interactions in an efficient and economical way. Methods 170, 38–47 (2020).

47. Lovell, J. T. et al. GENESPACE tracks regions of interest and gene copy number variation across multiple genomes. Elife 11, e78526 (2022).

48. Holst, F. et al. Helixer–*de novo* prediction of primary eukaryotic gene models combining deep learning and a hidden markov model. bioRxiv 10.1101/2023.02.06.527280 (2023).

49. Shumate, A. & Salzberg, S. L. Liftoff: accurate mapping of gene annotations. Bioinformatics 37, 1639–1643 (2021).

50. Haas, B. J. et al. Automated eukaryotic gene structure annotation using EVidenceModeler and the program to assemble spliced alignments. Genome Biol. 9, R7 (2008).

51. Ondov, B. D. et al. Mash: fast genome and metagenome distance estimation using MinHash. Genome Biol. 17, 132 (2016).

52. Jia, K. H. et al. SubPhaser: a robust allopolyploid subgenome phasing method based on subgenome-specific k-mers. New Phytol. 235, 801–809 (2022).

53. Sun, P. et al. WGDI: a user-friendly toolkit for evolutionary analyses of whole-genome duplications and ancestral karyotypes. Mol. Plant 15, 1841–1851 (2022).

54. Camacho, C., et al. BLAST+: architecture and applications. BMC Bioinformatics 10, 421 (2009).

55. Pertea, M. & Pertea, G. GFF utilities: GffRead and GffCompare. F1000Res. 9, ISCB Comm J-304 (2020).

56. Cosentino, S., Sriswasdi, S. & Iwasaki, W. SonicParanoid2: fast, accurate, and comprehensive orthology inference with machine learning and language models. Genome Biol. 25, 195 (2024).

57. Katoh, K. & Standley, D. M. MAFFT multiple sequence alignment software version 7: improvements in performance and usability. Mol. Biol. Evol. 30, 772–780 (2013).

58. Suyama, M., Torrents, D. & Bork, P. PAL2NAL: robust conversion of protein sequence alignments into the corresponding codon alignments. Nucleic Acids Res. 34, W609–W612 (2006).

59. Capella-Gutiérrez, S., Silla-Martínez, J. M. & Gabaldón, T. trimAl: a tool for automated alignment trimming in large-scale phylogenetic analyses. Bioinformatics 25, 1972–1973 (2009).

60. Nguyen, L. T., Schmidt, H. A., von Haeseler, A. & Minh, B. Q. IQ-TREE: a fast and effective stochastic algorithm for estimating maximum-likelihood phylogenies. Mol. Biol. Evol. 32, 268–274 (2015).

61. Letunic, I. & Bork, P. Interactive Tree of Life (iTOL) v6: recent updates to the phylogenetic tree display and annotation tool. Nucleic Acids Res. 52, W78–W82 (2024).

62. Zhang, C., Rabiee, M., Sayyari, E. & Mirarab, S. ASTRAL-III: polynomial time species tree reconstruction from partially resolved gene trees. BMC Bioinformatics 19, 153 (2018).

63. Junier, T. & Zdobnov, E. M. The Newick utilities: high-throughput phylogenetic tree processing in the UNIX shell. Bioinformatics 26, 1669–1670 (2010).

64. Sayyari, E. & Mirarab, S. Fast coalescent-based computation of local branch support from quartet frequencies. Mol. Biol. Evol. 33, 1654–1668 (2016).

65. Yang, Z. PAML 4: phylogenetic analysis by maximum likelihood. Mol. Biol. Evol. 24, 1586–1591 (2007).

66. Zhang, L. et al. Phylotranscriptomics resolves the phylogeny of Pooideae and uncovers factors for their adaptive evolution. Mol. Biol. Evol. 39, msac026 (2022).

67. Rhodes, J. A., Baños, H., Mitchell, J. D. & Allman, E. S. MSCquartets 1.0: quartet methods for species trees and networks under the multispecies coalescent model in R. Bioinformatics 37, 1766–1768 (2021).

68. Smith, S. A., Moore, M. J., Brown, J. W. & Yang, Y. Analysis of phylogenomic datasets reveals conflict, concordance, and gene duplications with examples from animals and plants. BMC Evol. Biol. 15, 150 (2015).

69. Martin, S. H. & Van Belleghem, S. M. Exploring evolutionary relationships across the genome using topology weighting. Genetics 206, 429–438 (2017).

70. Sayyari, E. & Mirarab, S. Testing for polytomies in phylogenetic species trees using quartet frequencies. Genes (Basel) 9, 132 (2018).

71. Allman, E. S., Baños, H. & Rhodes, J. A. NANUQ: a method for inferring species networks from gene trees under the coalescent model. *Algorithm*. Mol. Biol. 14, 24 (2019).

72. Sukumaran, J. & Holder, M. T. DendroPy: a Python library for phylogenetic computing. Bioinformatics 26, 1569–1571 (2010).

73. Chen, S. Ultrafast one-pass FASTQ data preprocessing, quality control, and deduplication using fastp. iMeta 2, e107 (2023).

74. Ewels, P., Magnusson, M., Lundin, S. & Käller, M. MultiQC: summarize analysis results for multiple tools and samples in a single report. Bioinformatics 32, 3047–3048 (2016).

75. Bray, N. L., Pimentel, H., Melsted, P. & Pachter, L. Near-optimal probabilistic RNA-seq quantification. Nat. Biotechnol. 34, 525–527 (2016).

76. Majidian, S. et al. Orthology inference at scale with FastOMA. Nat. Methods 22, 269–272 (2025).

77. Li, H. Aligning sequence reads, clone sequences and assembly contigs with BWA-MEM. arXiv 10.48550/arXiv.1303.3997 (2013).

78. Poplin, R. et al. A universal SNP and small-indel variant caller using deep neural networks. Nat. Biotechnol. 36, 983 (2018).

79. Yun, T. et al. Accurate, scalable cohort variant calls using DeepVariant and GLnexus. Bioinformatics 36, 5582–5589 (2020).

